# cGAS-mediated IFN-I signaling contributes to disease progression in drug-refractory epilepsy

**DOI:** 10.64898/2026.01.30.702860

**Authors:** Yige Huang, Li Fan, Man Ying Wong, Zhuofan Lei, Balaji Krishnamachary, Daphne Zhu, Mika Cadiz, Ravi Kumar Nagiri, Pearly Ye, Kendra Norman, Maitreyee Bhagwat, Young Jae Lee, Hui Li, Jingjie Zhu, Sadaf Amin, Kelli Lauderdale, Hao Chen, Wenjie Luo, Shiaoching Gong, Benjamin L. Liechty, Jorge J. Palop, Subhash C. Sinha, Junfang Wu, Mingrui Zhao, Li Gan

## Abstract

Epilepsy is a prevalent neurological disease with a third of patients becoming non-responsive to antiepileptic drugs and developing drug-refractory epilepsy (DRE). Here we report that DRE disease progression is contributed by overactive cyclic GMP-AMP synthase (cGAS), a double-stranded DNA sensor that induces type I interferon (IFN-I) signaling. In human DRE microglia, we observe a robust IFN-I signature and the activation of upstream cGAS-STING signaling. Further, in mouse models of Dravet syndrome, a genetic form of DRE, we observe the activation of the cGAS pathway. We show that microglial cGAS can be activated by DNA released from hyperexcitable neurons. Genetic reduction and pharmacological inhibition of cGAS reduces epileptic phenotypes, glial inflammatory signatures, and neuronal transcriptomic changes, underscoring the therapeutic potential of targeting cGAS for DRE treatment.

## INTRODUCTION

Epilepsy is a prevalent neurological disorder affecting approximately 65 million people worldwide, spanning all age groups^1^. For most patients, the primary treatment involves antiepileptic drugs (AEDs). Despite over 20 available AEDs, one-third of adult patients and about 20-25% of children with epilepsy experience uncontrolled seizures, defining them as drug-resistant epilepsy (DRE) cases if they fail to achieve seizure freedom after trials of two AEDs^2^. DRE is notably common in genetic forms of epilepsy, such as Dravet syndrome (DS), a severe developmental and epileptic encephalopathy that begins in infancy^3^. Over 80% of DS cases are linked to variants in the SCN1A gene, which codes for the voltage-gated sodium channel Nav1.1, leading to a haploinsufficiency^4^. DS patients often suffer from spontaneous seizures, developmental delays, autism spectrum disorder characteristics, and an elevated risk of sudden unexpected death in epilepsy (SUDEP)^5, 6^. Recently, the FDA approved three new anticonvulsants, but the inherent drug resistance of DS underscores the urgency to explore alternative therapeutic approaches^7^.

Epilepsy arises from various etiologies beyond genetic mutations, including structural, infectious, metabolic, and immune factors^8^. Neuroinflammation is a unifying feature across these conditions. Elevated pro-inflammatory cytokines and reactive astrocytes and microglia are frequently observed in drug-resistant epileptogenic brain tissues and animal models, implicating cytokines such as IL-1β, TNF-α, and IL-6 in seizure promotion^9, 10^.

Neuroinflammation in epilepsy can be initiated by the innate immune response to various pathogen-associated molecular patterns (PAMPs) and damage-associated molecular patterns (DAMPs)^11–14^. DNA is one type of DAMP linked to epileptogenesis, with mitochondrial DNA (mtDNA) damage, defective DNA repair, and activation of DNA-sensing toll-like receptors (TLRs) implicated in seizure development^15–17^. The cyclic GMP-AMP synthase (cGAS) is a crucial type of DNA sensor for cytosolic DNA and activates the stimulator of interferon genes (STING) to induce type I interferon (IFN-I) expression through the TBK1 signaling pathway^18, 19^. Previous work showed that cGAS-STING-IFN-I activation disrupts neuronal activity by inhibiting the MEF2C transcriptional network, a key regulator of neuronal function, where MEF2C haploinsufficiency is known to cause seizures^20, 21^. These findings highlight the role of the cGAS-mediated DNA sensing pathway in regulating neuronal activity and network function.

In this study, we investigated the involvement of the cGAS-STING-IFN-I pathway in DRE. We identified a strong IFN-I signature in the microglia of human DRE patients from single-cell transcriptomic data. Using DS as a DRE model, we showed that cGAS reduction and inhibition alleviated epileptic phenotypes, reduced inflammation, and normalized transcriptional changes in neurons and glia. These results highlight the potential of targeting cGAS as a therapeutic strategy for DS and other forms of DRE.

## RESULTS

### cGAS-STING-IFN-I is elevated in human DRE brains

Microglia, the resident immune cells in the brain, are known to modulate neuronal activity by suppressing neuronal activation, and the loss of microglia homeostatic receptor P2RY12 can promote seizures^22, 23^. We first analyzed a published single-cell transcriptomic dataset of immune cells from 11 brain tissues (hippocampal and cortical) from six individual patients (4 children and 2 adults) with DRE, which identified profound pro-inflammatory signatures^24^. Fifteen microglia subclusters were identified among all immune cells (**Fig. 1a**). Expression of interferon-stimulated genes (ISGs) (*STAT1, PARP14, RNF213, IFIT3*, and others) were enriched in clusters 3 and 4, but highest in cluster 4 (**Fig. 1b**). Genes associated with interferon-alpha and gamma response, as well as transcription factor (TF) motifs of IFN response factors (IRFs) and IFN-sensitive response elements (ISREs) were enriched in the marker genes of cluster 4 (**Fig. 1c, d**, table S1). Using ingenuity pathway analysis (IPA), cGAS and STING were identified as potential upstream regulators that led to the IFN response in cluster 4 microglia (**Fig.1e**, table S1).

**Figure. 1:**
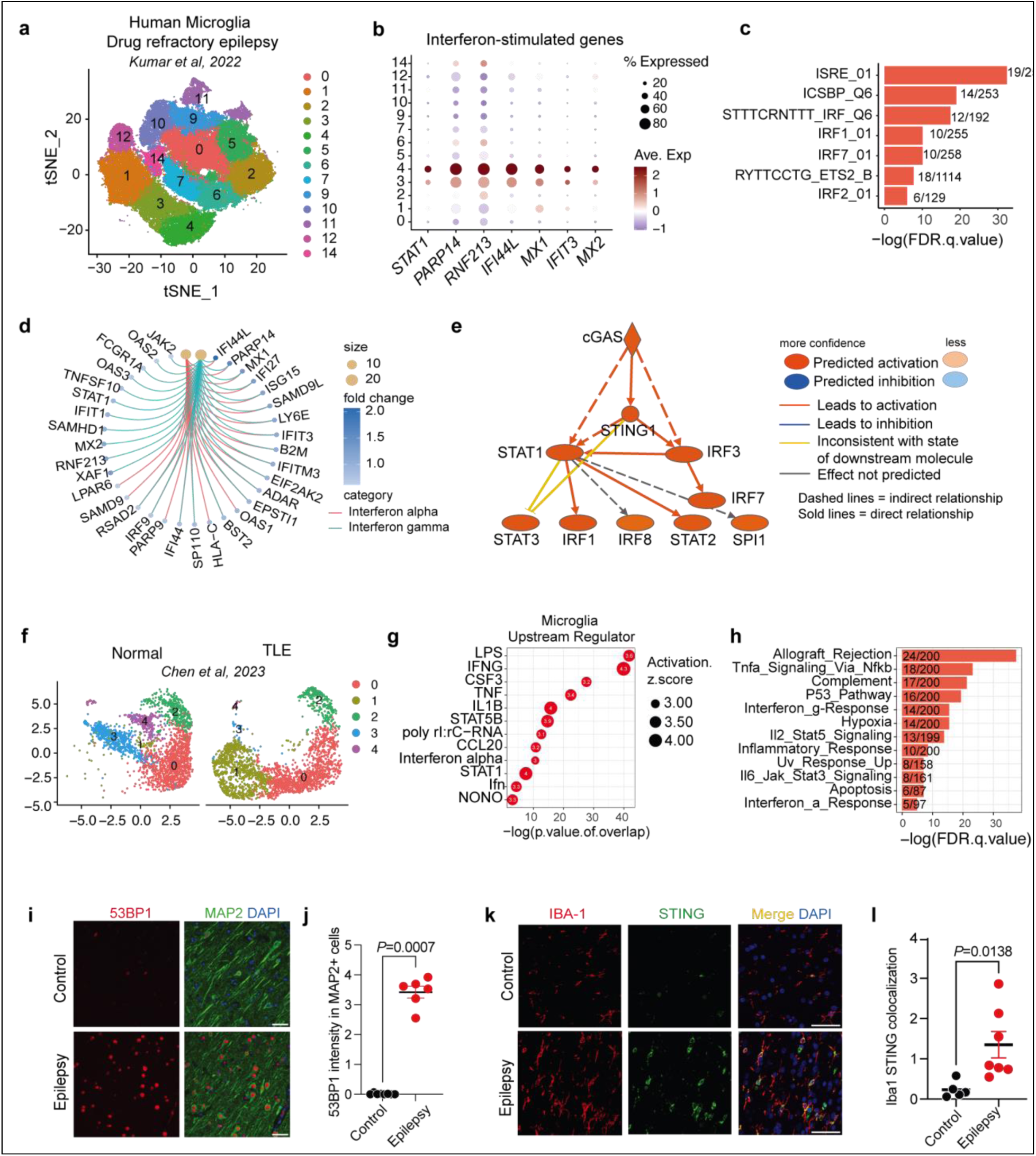
cGAS-STING-IFN-I is elevated in human DRE brains. (a) UMAP plot of microglia clusters sequenced from brain tissue obtained during resective epileptic brain surgery reported by Kumar et al, 2022^24^. (b) Dotplot showing the expression of interferon-stimulated genes (ISGs) across microglia clusters. c) Gene set enrichment analysis (GSEA) showing the top transcription factors (TF) associated with the marker genes of microglia cluster 4. (d) Cnetplot showing the logFC of microglia cluster 4 marker genes associated with the interferon pathways. (e) IPA predicted network showing the activation of cGAS and its downstream targets. IPA predicts cGAS as an upstream regulator of IFN-high cluster 4 of human DRE microglia (activation z-score = 2.23, *P* < 0.0001). (f) UMAP plot of microglia clusters split by disease condition sequenced from freshly dissected cortex samples of TLE participants and age-matched non-TLE control (n=4 each) reported by Chen et al, 2023^25^. (g) Dot plot showing top IPA upstream regulator predictions based on upregulated DEGs comparing TLE versus control participants. (h) GSEA analysis showing top hallmark pathways associated with upregulated DEGs comparing TLE versus control participants. (i) Representative 20x immunofluorescence images of 53BP1 and MAP2 in the human control and epilepsy brain cortical sections. Scale bar, 50um. (j) Quantification of 53BP1 intensity in MAP2^+^cells over the area of the image. Each dot represents the mean quantification of five images per section. Each section represents one case. Mann-Whitney test. Control, n=8; Epilepsy, n=6. (k) Representative image and (l) quantification of IBA-1 and STING colocalization in human control (n=5) and epilepsy (n=7) cortices. Each dot represents the average intensity of 4-5 images per section. Each section represents one case. Scale bar, 50um. Welch’s t-test.

Analysis of another published single-nuclei dataset from both drug-refractory temporal lobe epilepsy (TLE) patients and normal controls^25^ identified a shift in microglia clusters comparing TLE to normal controls (**Fig. 1f**). IPA upstream regulator analysis and GSEA hallmark analysis on the upregulated differentially expressed genes (DEGs) in TLE microglia identified IFN-I signaling to be a top player in the microglial response to epileptic lesions (**Fig. 1g, h**, table S2).

Considering the role of cGAS as a sensor for double-stranded DNA (dsDNA), we hypothesized that repetitive seizures might cause DNA damage in neurons and DNA release from degenerating neurons, which microglia could engulf^16, 26^, leading to cGAS-STING activation. Supporting this hypothesis, immunostaining revealed a significant increase in 53BP1, a DNA damage marker, in cortical neurons from epileptic lesions in DRE patients compared to controls (**Fig. 1i, j**, table S3). Further, to confirm the activation of cGAS-STING-IFN pathway in microglia of epilepsy patients, we performed immunofluorescent staining of STING and the microglia marker IBA-1. Indeed, microglial STING expression was increased in the cortical regions of human epilepsy brains compared to the normal control (**Fig. 1k, l**), suggesting the involvement of cGAS-STING activation in promoting microglial IFN response associated with epilepsy in human.

### Neuronal hyperexcitability activates microglial cGAS in vivo and in vitro

*SCN1A* haploinsufficiency-induced DS is a well-established form of DRE in need of novel therapeutic targets due to its devastating and lifelong nature. Thus, we were interested in understanding if maladaptive responses in microglia could contribute to the disease progression in DS. We first performed snRNAseq of both frontal cortex and hippocampal tissue of 6-month-old conditional Dravet (*PV-cre/+, Scn1a F/+*) and littermate control (*PV-cre/+, Scn1a +/+*) mice. 86,698 nuclei passed quality control and were clustered into different cell types based on the expression of specific marker genes (**Extended Data Fig. 1**). Microglia, with other immune cells, marked by the expression of *Cx3cr1*, *P2ry12*, and *Csf1r*, were further subclustered. Macrophage and T cell groups were removed based on the expression of *Mrc1* and *Skap1*, respectively. Final microglia cells were grouped into 5 distinct clusters with cluster shifts observed in the hippocampi but not the cortices of these conditional Dravet mice (**Fig. 2c**). Specifically, a significant loss of microglia cluster 1 and an expansion of microglia cluster 2 were observed (**Fig. 2d-e**). Pseudo-bulk analysis of DEGs showed upregulation of genes associated with the antigen presentation (*H2-K1, H2-D1, Nlrc5*) and interferon response (*Stat1, Sp100*) (**Fig. 2a, b**, table S4). Using SCENIC^27^ to predict transcription factor activity in different microglia clusters, oxidative stress and interferon-related TFs such as *Nfe2l2*, *Stat1*, and *Stat2* were identified to be highly active in cluster2 (**Fig. 2f**). Additional analyses using high-dimensional weighted correlation network analysis (hdWGCNA)^28^ revealed three modules of highly correlated genes: turquoise, brown, and blue. Specifically, the turquoise module was mostly expressed by cluster 2 microglia, and the blue module was mostly expressed by cluster 1 (**Fig. 2g**). Hallmark analysis of feature genes of each module suggested the turquoise module was enriched for interferon-gamma and alpha response, while the blue module was enriched for pathways important for homeostatic microglia function such as TGF-beta signaling (**Fig. 2g, Extended Data Fig. 2a–c**, table S4).

**Figure. 2:**
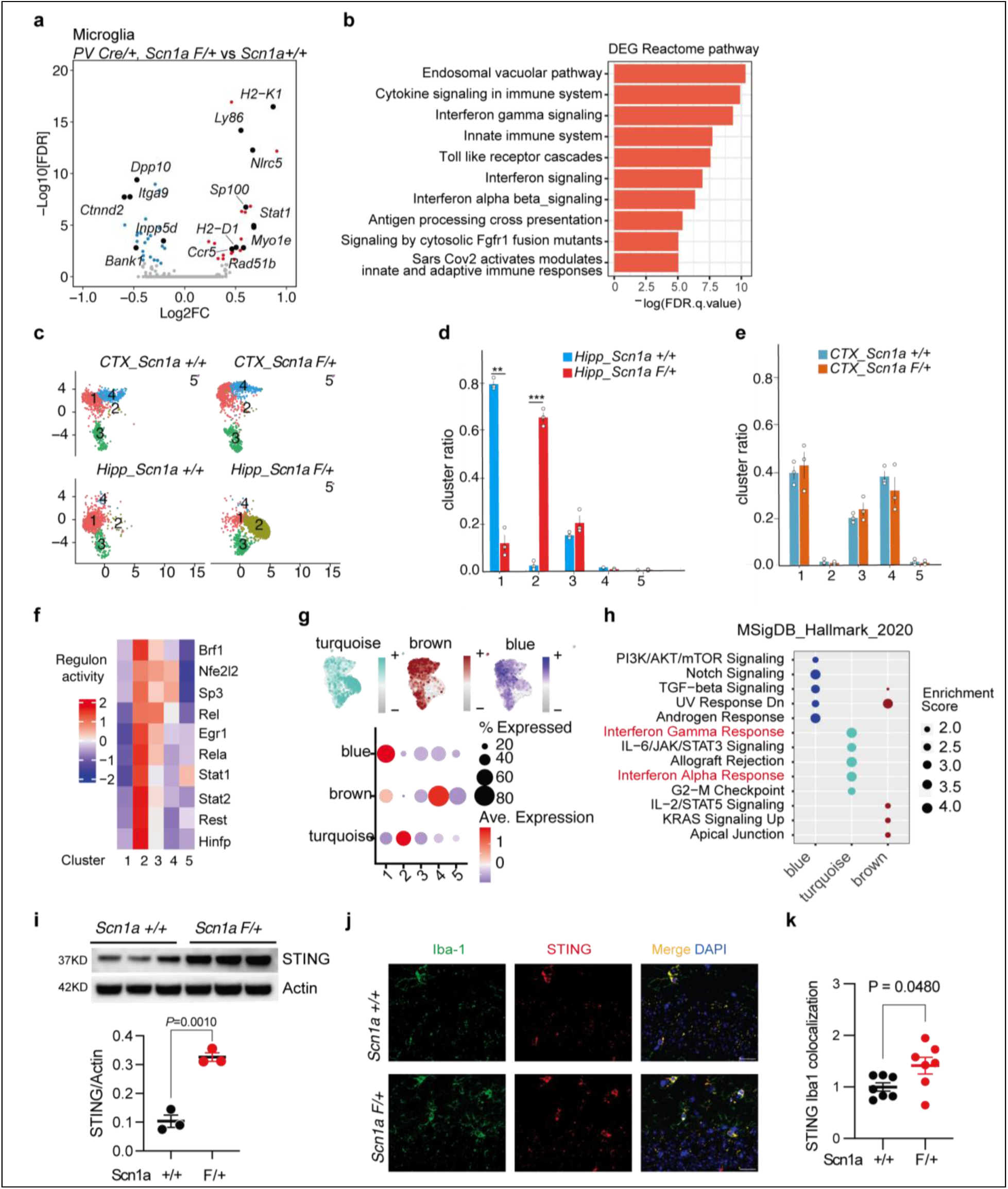
Neuronal hyperexcitability activates microglial cGAS-STING in *vivo* and in *vitro*. (a) Volcano plot of the DEGs comparing *PV-cre/+, Scn1a F/+* versus *PV-cre/+, Scn1a+/+* microglia. Red and blue dots represent significant DEGs ((logFC >=0.1 or <= -0.1, *p*.adj <0.05). (b) Bar plot of top pathways associated with upregulated DEG in *PV-cre/+, Scn1a F/+* microglia identified by GSEA hallmark analysis. (c) UMAP plot of microglia clusters split by genotype and brain region from *PV-cre/+, Scn1a +/+* and *PV-cre/+, Scn1a +/-* mice (n=3/genotype). (d-e) Bar plot of the ratio of microglia that belongs to each cluster in the hippocampus (d) and (e) cortex region of *PV-cre/+, Scn1a +/+* and *PV-cre/+, Scn1a +/-* mice. Each circle represents an individual animal. \*\**P* = 0.004, \*\*\**P* = 0.0007. Two-way ANOVA analysis with Sidak’s multiple comparison. (f) Heatmap of the selected transcription factor regulon activity across microglia clusters predicted by SCENIC. (g) Top: Feature plot showing the microglia cells that belong to each module predicted by hdWGCNA in UMAP. Bottom: Dotplot of the expression of each hdWGCNA module across all microglia clusters. (h) GSEA hallmark analysis showing the top pathways associated with each hdWGCNA module. (i) Western blot image and quantification of STING normalized to β-Actin in *PV-cre/+, Scn1a +/+* mice (n=3) and *PV-cre/+, Scn1a F/+* mice (n=3) hippocampal lysate. Unpaired t-test. (j) Representative immunofluorescence images of Iba1 and STING in the hippocampus of PV-cre/+, Scn1a F/+ mice and control mice. Scale bar, 20µm. (k) Quantification of STING IBA1 colocalization within the hippocampus. Each circle represents the mean quantification of four brain sections per animal. PV-cre/+, Scn1a +/+, n=7; PV-cre/+, Scn1a F/+ mice, n=7. Welch’s t-test.

Consistent with the regional specificity, immunoblotting revealed significantly increased STING expression in the hippocampus, not cortex, of *PV-Cre/+, Scn1a F/+* mice compared with littermate controls (**Fig. 2i**). Similarly, immunofluorescence analysis of STING and IBA1 demonstrated increased STING colocalization within hippocampal microglia, but not cortical microglia, in *PV-Cre/+*, *Scn1a F/+* mice (**Fig.2j-k, Extended Data Fig. 2e–f**), supporting activation of the microglial cGAS–STING pathway in this Dravet syndrome model.

### Neuronal hyperexcitability activates microglial cGAS–STING via DNA release

cGAS-STING signaling is initiated by cGAS sensing cytosolic dsDNA. To assess how this mechanism is engaged in DS mice, we quantified extranuclear dsDNA using a combined immunohistochemistry and EV profiling approach (**Fig. 3a**). Brain sections were stained for DAPI, Iba1, and dsDNA with outlined sampling windows within hippocampal CA1 (**Fig. 3b**), followed by high magnification analysis of individual microglia (**Fig. 3c**). Notably, the *Scn1a +/-* mice exhibited a significant accumulation of dsDNA puncta in the cytoplasm of hippocampal microglia (**Fig. 3c d**), supporting a microglia specific increase in cytosolic dsDNA associated with epileptic activity.

**Figure. 3.**
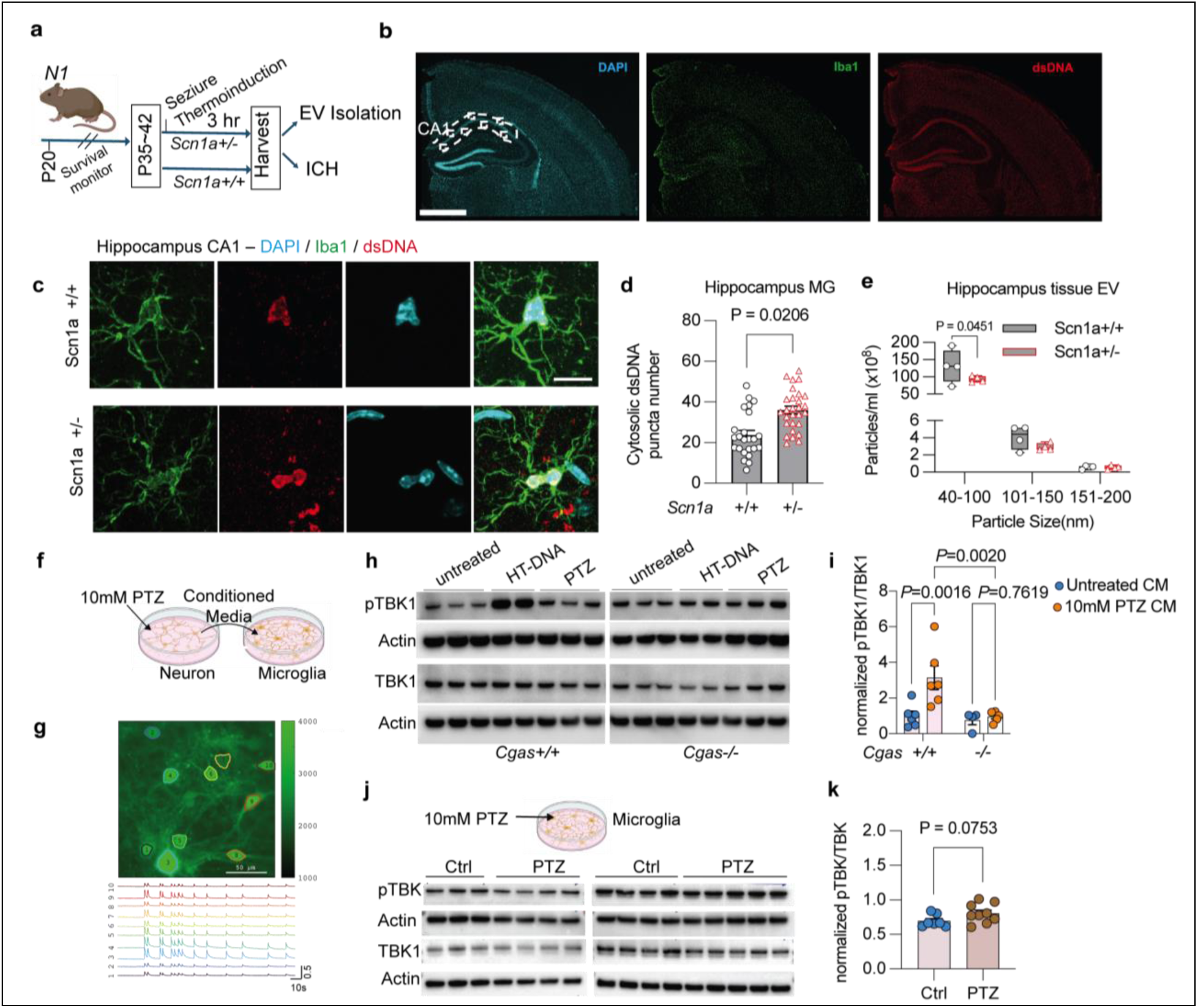
Hyperexcitability elevates dsDNA in microglia and activates cGAS-STING pathway. (a) Schematic showing the experiment design of the EV isolation and ICH in Dravet mice. (b) Confocal image of the brain section stained with DAPI, Iba1, and double-strain DNA (dsDNA), displaying sampling windows in hippocampus CA1. Bar = 1000 ⎧m. (c) 100x images from *Scn1a +/+* and *Scn1a +/-* mouse showing immuno-fluorescence signals of DAPI, Iba1, and dsDNA in hippocampus CA1 single microglia cells. Bar=10 ⎧m. (d) Analysis of total dsDNA puncta in the cytoplasma of single hippocampal microglia cells and comparison between the two groups. N=79 cells (*Scn1a +/+*), 101 cells (*Scn1a +/-*). N = 4 mice per group. Mixed-effects model ANOVA (F(1,6)= 9.742). (e) NanoFCM showed significant changes in size-based distribution in of hippocampal tissue EVs. N = 4 mice per group, two-way ANOVA (F (2, 18) = 78.95), Sidak’s multiple comparisons test. (f) Experimental scheme for the *in vitro* identification of the endogenous neuronal ligand for cGAS activation in primary microglia. (g) Acute PTZ-induced spontaneous epileptic activity in primary neurons. Up: representative fluorescence images of neurons expressing GCaMP8f (up). Scale bar: 50 μm. Down: calcium times course of each cell from above image showed spontaneous epileptic events after treated with 10 mM PTZ. (h) Representative western blot image for pTBK1, TBK1, and β-actin in primary *Cgas+/+* and *Cgas-/-* microglia treated with untreated neuronal CM, HT-DNA, and PTZ-treated neuronal CM. (i) Quantification of the ratio of pTBK1 to total TBK1 protein level. Each dot represents one well from two biologically independent experiments. *Cgas+/+*: untreated CM, n=6, PTZ-treated CM, n=6. *Cgas-/-*: untreated CM, n=4, PTZ-treated CM, n=5. Two-way ANOVA with uncorrected Fisher’s LSD. (j) Experimental scheme for the *in vitro* identification of the endogenous neuronal ligand in PTZ-treated primary microglia (up). Representative western blot image for pTBK1, TBK1, and β-actin in primary *wt* microglia treated with control (ctrl) and PTZ (bottom). (k) Quantification of the ratio of pTBK1 to total TBK1 protein level. Each dot represents one well from two biologically independent experiments. *Ctrl*, n=7, PTZ, n=9. Welch’s t test.

Because EVs can carry dsDNA^29–31^, we next examined tissue-derived EVs using Nano-flow cytometry measurement (NanoFCM). Scn1a+/− mice exhibited a selective reduction in small EVs within the exosome size range of approximately 40 to 100 nm in the hippocampus (**Fig. 3e**). Small EVs have been implicated in the export of nuclear and mitochondrial dsDNA from cells, and their reduction is therefore consistent with reduced dsDNA clearance and elevated cytosolic accumulation in microglia.

To directly assess whether neuronal hyperexcitability could trigger microglial cGAS activation, we turn to in vitro cell culture system (**Fig. 3f**). Hyperexcitability of neurons were triggered with 10 mM pentylenetetrazole (PTZ), which disrupts inhibitory synaptic transmission mediated by GABA-A receptors. Using a genetically encoded calcium sensor GCaMP8-fast transduced with a lentivral vector (Lenti-hSynapsin-jGCaMP8f), we confirmed spontaneous epileptiform activities after acute perfusion with 10 mM PTZ, as evidenced by synchronized high-amplitude ictal-like events and high-amplitude interictal spikes (**Fig. 3g**). After 24 hr, we collected conditioned media (CM) from PTZ-treated or non-treated primary neurons and applied it to primary microglia with or without cGAS (**Fig. 3f**). Western blot analysis showed that both Herring testis DNA (HT)-DNA, serving as a positive control, and CM from PTZ-treated neurons increased TBK1 phosphorylation in *Cgas^+/+^* microglia compared to CM from untreated neurons. Deleting cGAS abolished the p-TBK elevation induced by CM from PTZ-treated neurons (**Fig. 3h-i**). To exclude the possibility that PTZ directly affects microglial pTBK1, primary microglia were treated with 10mM PTZ or control media for 1 h, followed by microglia collection 24 hrs later (**Fig. 3j**). Western blot analysis revealed no difference in TBK1 phosphorylation between control and PTZ-treated group (**Fig. 3k**). Taken together, our findings suggest that DNA released from hyperexcitable neurons can activate cGAS-STING in microglia.

### cGAS deletion ameliorates pharmacologically-induced seizures

Our results so far provide strongly evidence of microglial activation of cGAS-STING in the presence of seizure-associated hyperexcitability. We next employed the acute PTZ model to assess the contribution of cGAS to seizure susceptibility under conditions of transient network disinhibition. *Cgas^−/−^* mice and littermate *Cgas^+/+^*controls were treated with PTZ (50 mg/kg, i.p), with a second challenge performed three weeks later to assess the effects on seizure sensitization (**Fig. 4a**). Seizure severity was quantified using the modified Racine scale^32^, recording the maximum score during each session. Male *Cgas^−/−^* mice consistently exhibited lower seizure scores than *Cgas^+/+^* littermates in both PTZ treatments (**Fig. 4b**). Female *Cgas^−/−^* mice showed a trend toward reduced seizure scores during the first challenge and a significantly increased latency to tonic clonic seizures during the second challenge (**Fig. 4c d**).

**Figure. 4:**
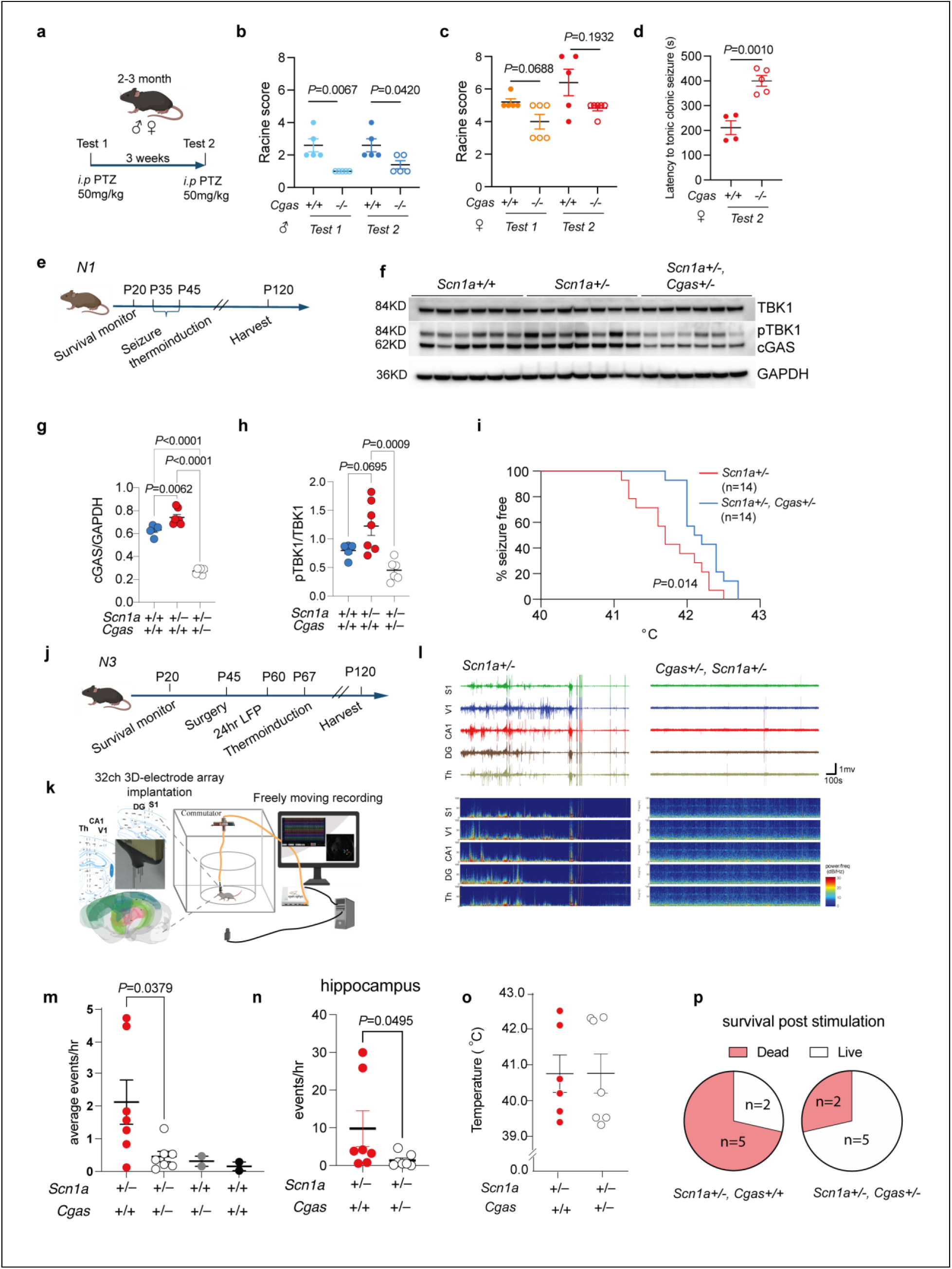
cGAS knockdown and inhibition ameliorates pharmacological and Dravet-associated seizure phenotypes. (a) Schematic showing the experiment design of the pharmacological model of seizures using PTZ. (b) Maximum Racine score of seizures induced by PTZ in male *Cgas+/+* mice (n=5) and *Cgas-/-* mice (n=6). Mann-Whitney test. (c) Left: Maximum Racine score of seizures induced by PTZ in female *Cgas+/+* mice (n=5) and *Cgas-/-* mice (n=5). Mann-Whitney test. (d) Latency to tonic-clonic seizure in seconds in mice with Racine score >=5. Unpaired t-test. (e) Schematic showing the experiments performed in N1 mice along the timeline. (f) Western blot image and (g, h) quantification of cGAS protein level normalized to GAPDH and pTBK1 protein level normalized to total TBK1 in the hippocampal lysate of N1 *Scn1a+/+* (n=5, after one significant outlier removed), *Scn1a+/-* (n=7), and *Scn1a+/-, Cgas+/-* (n=6) mice. Ordinary one-way ANOVA with Tukey’s multiple comparison test. (i) Quantification of seizure-free temperature in N1 mice after thermal induction. *Scn1a+/-, Cgas+/+*: n=14, 6 male, 8 female; *Scn1a+/-, Cgas+/-*: n=14, 7 male, 7 female. Log-rank Mantel-Cox test. (j) Schematic showing the experiments performed in N3 mice along the timeline. (k) Schematic showing the electrode implantation and experiment setup for 24hr LFP recording. (l) Representative LFP traces (top) and spectrogram(bottom) in N3 mice. (m) Quantification of the average number of epileptiform events per hour across all brain regions recorded in N3 *Scn1a+/-*(n=7), *Scn1a+/-, Cgas+/-* (n=7), *Scn1a+/+* (n=2), *Scn1a+/+, Cgas+/-*(n=2) mice. Mann-Whitney test used to compare *Scn1a+/-* and *Scn1a+/-, Cgas+/-*. (n) Quantification of the number of epileptiform events split by spike load in the hippocampal region in *Scn1a+/-* and *Scn1a+/-, Cgas+/-* mice. Mann-Whitney test. (o) Quantification of the seizure-free temperature in N3 mice after thermal induction. Each symbol denotes one animal. *Scn1a+/-, Cgas+/+* (n=6, one dead before thermal induction experiment); *Scn1a+/-, Cgas+/-* (n=7). (p) Pie chart showing the percentage breakdown of survival rate post stimulation from 24-hr recording and hyperthermia-induced seizures. *Scn1a+/-, Cgas+/+* (n=7); *Scn1a+/-, Cgas+/-* (n=7).

To investigate the microglia-specific cGAS function, *Cx3cr1^CreER^*mice were crossed with *cGas^tm^*^1c^ mice to generate *cGas^tm^*^1c^*: F/F; CX3CR1^CreER^ Cre/+* and littermate *cGas^tm^*^1c^ *F/F; CX3CR1^CreER^ +/+* control mice. Cre recombinase expression was induced through tamoxifen administration at the 2 months of age. We assessed the susceptibility to seizures in both groups by administering PTZ (50mg/kg, i.p.) at 19-23 months of age. *cGas^tm^*^1c^*: F/F; CX3CR1^CreER^ Cre/+* mice showed significantly lower maximum seizure scores compared to their control littermates (**Extended Data Fig. 4a**).

Our analysis of human temporal lobe epilepsy snRNA-seq data revealed robust activation of type I interferon signaling in microglia (Fig. 1f–h). We therefore tested whether cGAS deletion reduces seizure severity in the kainic acid model, a limbic excitotoxic paradigm that more closely reflects temporal lobe epilepsy^22, 23^. *Cgas^−/−^*and *Cgas^+/+^* mice were administered with intraperitoneal kainic acid (KA, 18 mg/kg) at 2-6 months of age. Consistent with the PTZ model, male *Cgas^−/−^*mice displayed significantly reduced seizure scores compared with *Cgas^+/+^* controls, while female *Cgas^−/−^*mice showed a similar trend (**Extended Data Fig. 4b, c**). These findings support a broad role for cGAS in promoting epileptogenesis.

### Genetic cGAS reduction ameliorates Dravet-associated seizure phenotypes

Building on the evidence of cGAS promotes PTZ and KA-induced seizures, and the elevation of cGAS–STING signaling in Dravet mice, we next examined if cGAS contributes to disease progression in DS. *Scn1a^tm1Kea^*mice on a pure 129S6/SvEvTac background (*Scn1a+/-129S6*) do not have overt Dravet phenotypes, but when backcrossed with C57BL/6J (*B6*) mice and on the N1 strain, exhibit spontaneous seizures, early lethality and susceptibility to febrile seizure^33^. We crossed *Scn1a+/-129S6* mice with *Cgas+/-B6* mice to generate N1 *Scn1a+/+* and *Scn1a+/-* mice that are *Cgas+/+* or *Cgas+/-* (**Extended Data Fig. 3a**).

We began the monitoring of survival at P20, assessed the hyperthermia-induced seizure susceptibility by raising body temperature at P35–P45, and harvested the surviving mice at around P120 (**Fig. 4e**). *Scn1a+/-* increased cGAS protein levels and showed a trend of elevated TBK1 phosphorylation, which was mitigated by *Cgas+/-* (**Fig. 4f-h**), confirming that the cGAS-STING pathway is activated in DS mice. Febrile seizure, a hallmark of DS, is commonly induced by infections and accompanied by inflammatory responses^34, 35^. To mimic febrile seizures, N1 mice were challenged by gradually raising body temperature with a heat lamp to a maximum of 42.5 °C or until clonic and tonic-clonic seizures occurred. N1 *Scn1a+/-Cgas+/-* mice exhibited a higher temperature threshold for seizures (mean = 42.2 °C) compared to *Scn1a+/-* littermates (mean = 41.7 °C) (**Fig. 4i**). However, no significant survival benefit was observed comparing N1 *Scn1a+/-, Cgas+/-* mice to the *Scn1a+/-* littermate **(Extended Data Fig. 3b**).

To assess whether cGAS reduction could mitigate the worsened epileptic phenotypes of *Scn1a+/-*mice on the B6 background^33, 36^, we backcrossed *Scn1a+/-* 129S6 mice with *Cgas-/-* or *Cgas+/+* B6 mice for three generations (**Extended Data Fig. 3c**) and performed experiments (**Fig. 4j**). Consistent with the observation in N1 *Scn1a+/-,* we observed no survival benefit for N3 *Scn1a+/-, Cgas+/-* mice compared to *Scn1a+/-* littermates (**Extended Data Fig. 3d**).

We next implanted 32-channel tetrodes into N3 DS brains at P60 and conducted 24-hour local field potential (LFP) recordings with video monitoring to capture spontaneous epileptiform activity across brain regions, including V1, S1, CA1, dentate gyrus, and thalamus (**Fig. 4k**). An automated algorithm was used to quantify epileptic spikes and identify epileptiform events with continuous spikes lasting over 10 seconds (**Fig. 4l**). *Scn1a+/-/Cgas+/-* mice exhibited fewer epileptiform events compared to *Scn1a+/-* littermates, especially in the hippocampi, suggesting reduced spontaneous spiking (**Fig. 4m, n**). N3 *Scn1a+/-, Cgas+/-* mice showed similar seizure thresholds to *Scn1a+/-* mice after hyperthermia challenges (**Fig. 4o**), they had reduced post-stimulation mortality, with 28.6% death observed until P120 compared to 71.4% in *Scn1a+/-*littermates (**Fig. 4p**). Taken together, genetic cGAS reduction ameliorates epileptic phenotypes in DS mice across genetic backgrounds and disease severities.

### Pharmacological inhibition of cGAS ameliorates Dravet-associated seizure phenotypes

To test the translational relevance of cGAS targeting in Dravet syndrome, we next evaluated whether pharmacological inhibition of cGAS mitigates DS-associated phenotypes in a cohort of N1 *Scn1a+/+* and *Scn1a+/-* mice by crossing *Scn1a+/-* 129S6 with wildtype B6. Starting at P15, the mice received daily intraperitoneal injections of 5 mg/kg of the brain-penetrant cGAS inhibitor TDI-6570^21^ (cGASi) or control until P21, followed by a daily, single dose of almond butter diet containing 5 mg/kg of cGASi or control. At P35, the mice were challenged with hyperthermia-induced seizures and harvested at P42 (**Fig. 5a**).

**Figure. 5:**
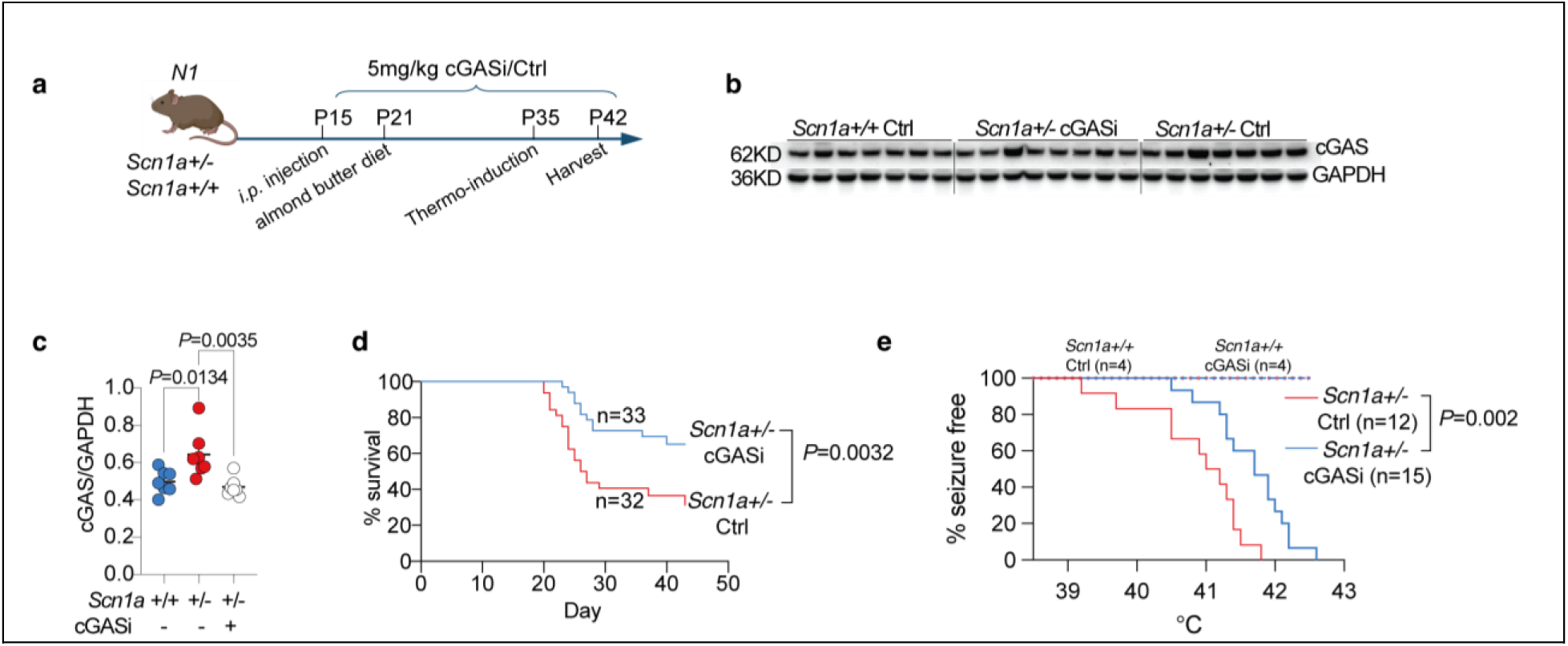
Pharmacological cGAS inhibition ameliorates Dravet-associated seizure phenotypes. (a) Schematic showing the treatment and experiments performed in N1 mice along the timeline. (b) Western blot image and (c) quantification of cGAS protein level normalized to GAPDH in the hippocampal lysate of N1 *Scn1a+/+,* Ctrl (n=7), *Scn1a+/-*, cGASi (n=8, one outlier removed with ROUT test, Q= 1% for quantification), *and Scn1a+/-*, Ctrl (n=7) mice. Ordinary one-way ANOVA with Tukey’s multiple comparison test. (d) Survival of *Scn1a+/-*, Ctrl (n=32) and *Scn1a+/-*, cGASi (n=33) monitored until harvest. Log-rank Mantel-Cox test. (e) Seizure-free ratio and temperature of *Scn1a+/+*, Ctrl (n=4), *Scn1a+/+*, cGASi (n=4), *Scn1a+/-*, Ctrl (n=12) and *Scn1a+/-*, cGASi (n=15) mice during hyperthermia-induced seizures. Log-rank Mantel-Cox test comparing *Scn1a+/-*, Ctrl and *Scn1a+/-*, cGASi mice.

*Scn1a+/-* control mice showed elevated cGAS expression compared to *Scn1a+/+* controls, which was prevented by cGASi treatment, indicating target engagement (**Fig. 5b, c**). cGASi treatment also reduced the IFN-I-induced chemokine CCL5 in *Scn1a+/-* mice (**Extended Data Fig. 5a**). Notably, cGASi markedly reduced early mortality in *Scn1a+/-* mice (**Fig. 5d**) and increased the mean hyperthermia-induced seizure threshold from 40.9°C to 41.7°C without affecting baseline body temperature (**Fig. 5e, Extended Data Fig. 5b**). None of the *Scn1a+/+* mice treated with control or cGASi experienced premature death or hyperthermia-induced seizures even at 42.5°C, demonstrating the therapeutic potential of cGAS inhibition for DS.

### Genetic cGAS reduction rescues DS-induced inflammatory signature in glial cells

We next interrogated cell type-specific transcriptomic changes induced by cGAS reduction in the *Scn1a+/-* mice. We focused on the hippocampus due to the strong inflammatory signatures induced by DS (**Fig. 1i, j**). SnRNAseq was performed in 4–5-month-old N3 *Scn1a+/+*, *Cgas+/-*; *Scn1a+/+*; *Scn1a+/-*; and *Scn1a+/-, Cgas+/-* mice (n=4/genotype). 137,028 nuclei were analyzed and grouped into different cell types after passing stringent QC (**Extended Data Fig. 6**). Among the 6 subclusters from 5,732 microglia nuclei, MG6 was mostly present in *Scn1a+/-* mice, while MG3 was almost exclusively present in *Scn1a+/-, Cgas+/-* mice (**Fig. 6a**). Expression of DAM genes (*Trem2, Ctsl, Cd9*), interferon receptor genes (*Ifnar2, Ifngr2*), antigen presentation genes (*H2-K1, H2-D1, Raet1e*) were highly enriched in *Scn1a+/-* and the associated cluster 6 microglia but dampened in *Scn1a+/-, Cgas+/-* and associated cluster 3 microglia **(Fig. 6b)**.

**Figure. 6:**
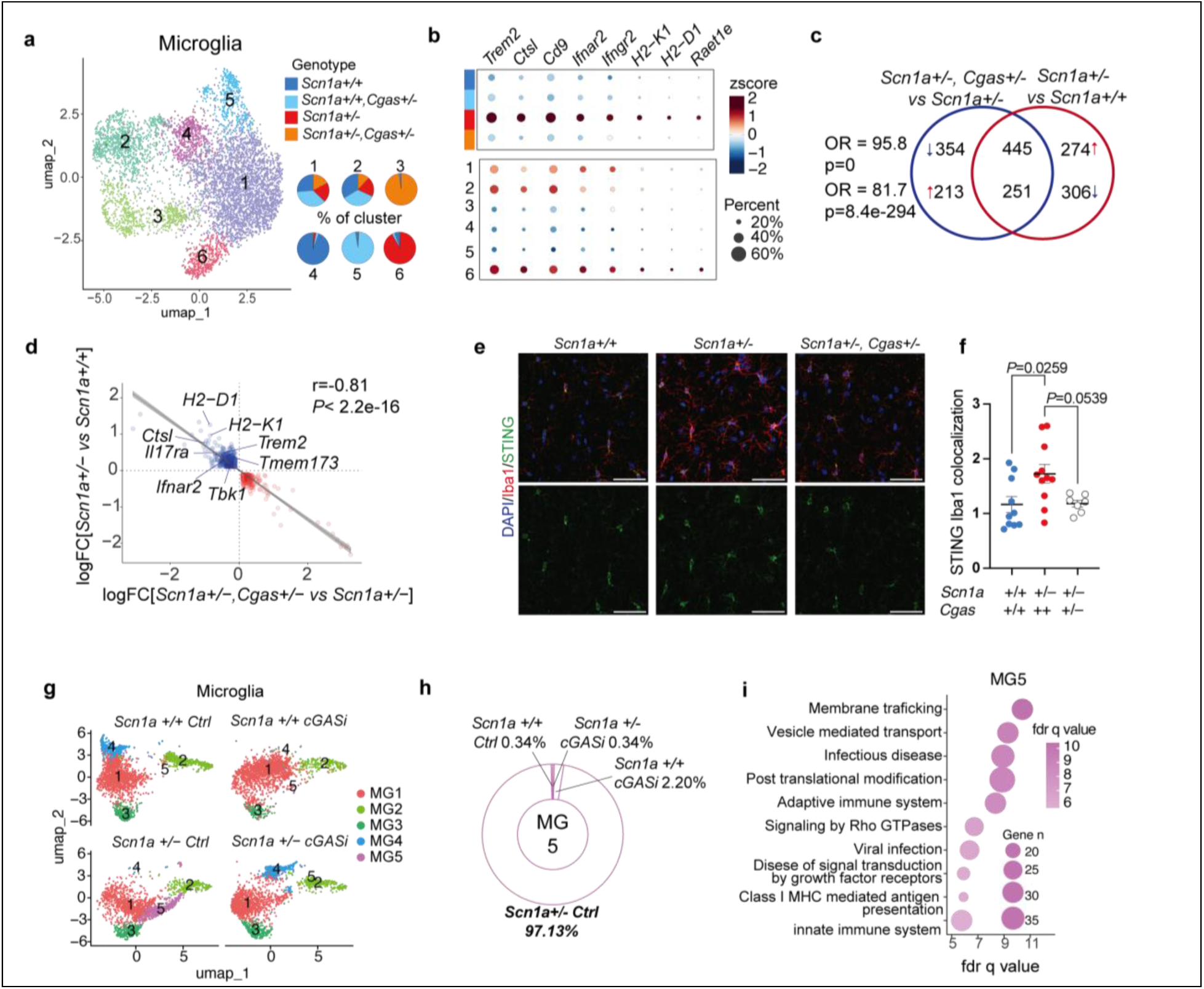
cGAS reduction and inhibition alleviates the inflammatory signature in Dravet microglia. (a) UMAP of microglia colored by clusters with a pie chart showing the percentage proportion of cells from different genotypes for each cluster. (n=2 per sex per genotype) (b) Dotplot showing the expression level and percentage expression of DAM genes (*Trem2, Ctsl, Cd9*), Interferon receptor genes (*Ifnar2, Ifngr2*), and antigen presentation genes (*H2-K1, H2-D1, Raet1e*) across genotypes and microglia clusters. (c) Venn diagram showing the overlap between the DEGs (logFC >=0.1 or <= -0.1, *p*.adj <0.05) of *Scn1a+/-, Cgas+/-* versus *Scn1a+/-*and *Scn1a+/-* versus *Scn1a +/+* microglia. Overlapping odds ratio and *P* value calculated with Fisher’s exact test using the GeneOverlap package^39^ in R. (d) Scatter plot and simple linear regression analysis with standard error showing a negative correlation between logFC values of overlapping DEGs in c). Pearson’s r = -0.81, *p*<2.2e-16. (e) Representative 20x immunofluorescence images of STING and IBA1 in the CA1 region of hippocampi of 4–5-month-old N3 mice. Scale bar, 50⎧m. (f) Quantification of STING IBA1 colocalization within the CA1 region. Each circle represents the mean quantification of three to four brain sections per animal. One-way ANOVA with Tukey’s multiple comparisons test. *Scn1a+/+, Cgas+/+*, n=10; *Scn1a+/-, Cgas+/+*, n=11; *Scn1a+/-, Cgas+/-*, n=7. (g) UMAP of microglia colored by clusters in hippocampal tissue of *Scn1a+/+,* Ctrl; *Scn1a+/+*, cGASi; *Scn1a+/-*, Ctrl; and *Scn1a+/-*, cGASi mice (n=4 per group). (h) Donut plot of the percentage genotype composition of microglia cluster 5 showing cluster 5 is mainly present in *Scn1a+/-,* Ctrl mice. (i) Dot plot of the GO pathways associated with microglia cluster 5 suggesting enrichment for infectious and immune response.

Pseudobulk analysis of DEGs revealed an upregulation of the above genes as well as members of the cGAS pathway including *Tmem173*(STING) and *Tbk1*, in *Scn1a+/-* microglia (**Extended Data Fig. 7a**, table S5). Indeed, cGAS reduction led to a significant rescue of 696 out of 1276 DEGs induced by *Scn1a+/-* (**Fig. 5c**, table S5). Linear correlation analysis showed a negative Pearson’s *r* of -0.81 between the log fold change of overlapping DEGs (*H2-D1, Tmem173, Tbk1,* and others) (**Fig. 6d**). GO analysis identified the top pathways modified by *Cgas+/-* were related to leukocyte migration/activation, inflammatory response, and phagocytosis (**Extended Data Fig. 7b, c**). hdWGCNA analysis identified 3 gene modules expressed by microglia. Using differential module eigengene (DME) analysis, we observed that *Scn1a+/-* induced but *Cgas+/-* reduced a turquoise gene module in microglia that was enriched in complement signaling, TLR signaling, and IFN-I response (**Extended Data Fig. 7d, e**). Immunostaining confirmed that STING expression in IBA-1+ microglia was elevated in the hippocampus of *Scn1a+/-* mice, but not when cGAS is genetically knocked down (**Fig. 6e, f**).

We further performed snRNAseq on N1 *Scn1a+/+* Ctrl, *Scn1a+/+* cGASi, *Scn1a+/-* Ctrl, and *Scn1a+/-* cGASi mice that received thermo-stimulation (n=4 per group). From 153,140 nuclei passing stringent QC, distinct cell types were identified (**Extended Data Fig. 8**). The microglia cells fall into five distinct subpopulations (**Fig. 6g**) In the control group, *Scn1a+/-* led to the emergence of cluster 5 microglia, which was absent in *Scn1a+/+* littermates. cGASi treatment in *Scn1a+/-* mice restored microglial clusters to a profile similar to *Scn1a+/+* controls (**Fig. 6h, Extended Data Fig. 7f**). GSEA of cluster 5 microglia revealed enrichment in immune-related pathways such as “infectious disease,” “adaptive immune system,” and “viral infection,” with high expression of genes including *Rnf130, Ctsb, Ube2k,* and *Ifngr2* (**Fig. 6i**, **Extended Data Fig. 7g**, Table S6). Together, microglial inflammatory response is activated in both acute and chronic stages of DS and can be markedly ameliorated by cGAS knockdown and inhibition.

Astrocytes are another important type of glial cell and have been reported to play crucial roles in the pathogenesis of epilepsy through regulating neurotransmitters, synaptic transmission, and inflammation^37^. In *PV-cre/+, Scn1a F/+* mice, we observed striking changes in astrocyte transcriptional states in the hippocampus, but not in the cortex (**Extended Data Fig. 9a, b**). Pseudobulk DEG analysis identified an upregulation of disease-associated astrocyte (DAA) markers^38^ such as *Gfap, Cd9, Id3, Aqp4* in *PV-cre/+, Scn1a F/+* astrocytes (**Extended Data Fig. 9c**, table S4). Correlation analysis of the cluster 4 marker genes and DAA marker genes showed a significant positive correlation with a Pearson’s *r* of 0.71, suggesting the expansion of DAA in the conditional Dravet mouse model (**Extended Data Fig. 9d**, table S4). Using western blot, we confirmed an increased expression of GFAP in the hippocampus of the conditional Dravet mice (**Extended Data Fig. 9e**).

Further analysis of the N3 *Scn1a+/-* snRNAseq dataset showed that among the 8 astrocyte clusters, cluster 4 was expanded in *Scn1a+/-* mice and rescued by *Cgas+/-* (**Extended Data Fig. 10a, b**). Confirming our observation in the conditional dataset, cluster 4 highly expressed DAA marker genes (**Extended Data Fig. 10c**). Pseudobulk analysis showed that *Cgas+/-* rescued 358 out of 549 *Scn1a+/-* induced astrocytes DEGs (**Extended Data Fig. 10d**, table S5). A negative correlation of *r* = -0.88 between overlapping DEGs was observed (**Extended Data Fig. 10e**). The elevated expression of DAA marker genes (*Gfap, Aqp4, Cd9, Clu, Apoe,* etc.) in *Scn1a+/-* were abolished by *Cgas* reduction (**Extended Data Fig. 10f**). GO analysis identified gliogenesis, glial cell differentiation, and regulation of protein catabolic process to be top pathways upregulated by *Scn1a+/-* and modified by *Cgas+/-* (**Extended Data Fig. 10f**). APOE^+^ lipid-accumulated reactive astrocytes have been reported to promote disease progression in epilepsy^25^. Notably, *Apoe* was a top DEG rescued by cGAS reduction. Indeed, immunostaining of APOE and GFAP revealed that both the immunointensity of GFAP and the colocalization between APOE and GFAP were increased in the CA1 region of the hippocampus of *Scn1a +/-* mice and reduced by *Cgas+/-*(**Extended Data Fig. 10h, i**). Together, cGAS reduction markedly ameliorated the inflammatory signature in microglia and astrocytes of DS mice.

### Genetic reduction of cGAS reduction rescues neuronal transcriptomic changes induced by Dravet mutation

DS is primarily caused by the variants in *SCN1A* which controls the excitability of neurons. To investigate the neuronal mechanisms involved, we first examined whether cGAS reduction could increase Nav1.1 expression, thus rescuing its haploinsufficiency. Immunoblotting showed no elevation of Nav1.1 protein levels with cGAS reduction, suggesting other indirect mechanisms (**Extended Data Fig. 3e, f**).

We further analyzed our snRNAseq data to understand neuronal changes involved with *Scn1a* mutation and cGAS manipulation. In the conditional Dravet snRNAseq dataset, subclustering of excitatory neurons (EN) and inhibitory neurons (IN) (**Extended Data Fig. 11a, b**) did not reveal any significant cluster changes ((**Extended Data Fig. 11c, d**). Pseudobulk DEG analysis revealed changes associated with genes involved in excitability and synaptic transmission (table S4). Specifically in EN, we observed the downregulation of *Pde10a*, inhibition of which leads to increased neuronal excitability and seizure activity^40^, *Npas4*, an activity-responsive immediate early gene involved in DNA damage repair^41^, which also regulates *Homer1* expression for homeostatic scaling post-seizures^42^, and *Mef2c*, which regulates synapse formation and we previously reported to be regulated by cGAS-IFN signaling^21, 43^(**Fig. 7a**). In IN, in addition to the downregulation of *Pvalb* and *Scn1a*, we also observed genes related to neurotransmitter receptors and release, and potassium channels (**Fig. 7b, c**, table S4).

**Figure. 7:**
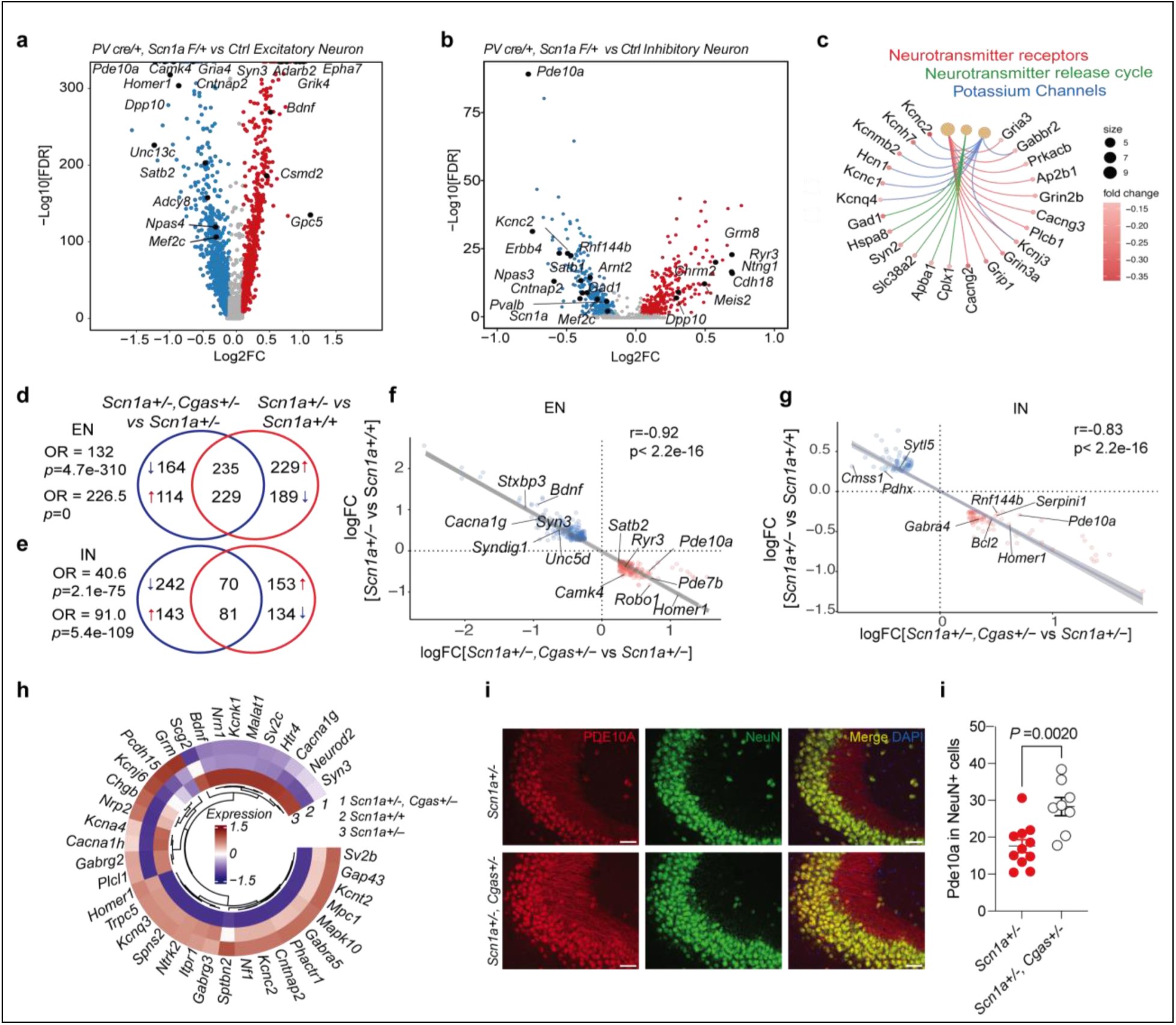
cGAS reduction rescues neuronal transcriptomic changes induced by Dravet mutation. **(a**) Volcano plot of the DEGs comparing *PV-cre/+, Scn1a F/+* versus *PV-cre/+, Scn1a +/+* excitatory neurons. Red and blue dots represent significant DEGs ((logFC >=0.1 or <= -0.1, *p*.adj <0.05). (b) Volcano plot of the DEGs comparing *PV-cre/+, Scn1a F/+* versus *PV-cre/+, Scn1a +/+* inhibitory neurons. Red and blue dots represent significant DEGs ((logFC >=0.1 or <= -0.1, *p*.adj <0.05). (c) Cnetplot showing the genes associated with the selected top GO pathway downregulated in *PV-cre/+, Scn1a F/+* versus *PV-cre/+, Scn1a +/+* inhibitory neurons. (d) Venn diagram showing the overlap between the top DEGs (logFC >=0.25 or <= -0.25, *p*.adj <0.05) of *Scn1a+/-, Cgas+/-* versus *Scn1a+/-* and *Scn1a+/-* versus *Scn1a+/+* excitatory neurons. Overlapping odds ratio and *P* value calculated with Fisher’s exact test. (e) Venn diagram showing the overlap between the top DEGs (logFC >=0.25 or <= -0.25, *p*.adj <0.05) of *Scn1a+/-, Cgas+/-* versus *Scn1a+/-* and *Scn1a+/-* versus *Scn1a +/+* inhibitory neurons. Overlapping odds ratio and *P* value calculated with Fisher’s exact test. (f) Scatter plot and simple linear regression analysis with standard error showing a negative correlation between logFC values of overlapping DEGs in D). Pearson’s r = -0.92, *p*<2.2e-16. (g) Scatter plot and simple linear regression analysis with standard error showing a negative correlation between logFC values of overlapping DEGs in E). Pearson’s r = -0.83, *p*<2.2e-16. (h) Heatmap showing the scaled average expression of seizure-related genes in *Scn1a+/+*, *Scn1a+/-*, and *Scn1a+/-, Cgas+/-* excitatory neurons. (i) Representative 20x immunofluorescence images of PDE10A and NeuN in the CA3 region of hippocampi of 4–5-month-old N3 mice. Scale bar, 50⎧m. (j) Quantification of PDE10A intensity within NeuN positive areas. Each circle represents the mean quantification of three to four brain sections per animal. Unpaired t-test. *Cgas+/+*, n=11; *Scn1a+/-, Cgas+/-*, n=8.

Genetic reduction of cGAS did not alter the subcluster composition of ENs or INs in the N3 *Scn1a+/−* cohorts (**Extended Data Fig. 11e-h**). Remarkably, *Cgas+/-* rescued 40% of the DEGs induced by *Scn1a+/-* in EN and around a third of DEGs in IN (**Fig. 7d, e**, table S5). Correlation analysis showed highly negative correlations between overlapping DEGs with r of -0.92 for EN and -0.83 for IN (**Fig. 7f, g**). We also observed common top DEGs with the conditional dataset which were rescued by *Cgas+/-*, including *Pde10a* and *Homer1* in both EN and IN and *Bdnf* ^44^ in EN, consistent with their crucial roles in epilepsy. Further, our snRNAseq data showed that *Cgas+/-* restored the expression of many seizure-related genes in *Scn1a+/-* EN back to *Scn1a+/+* levels (**Fig. 7h**). Using immunostaining, we validated that the expression of PDE10A in the CA3 neurons was upregulated in *Scn1a+/-, Cgas+/-* compared to *Scn1a+/-* mice (**Fig. 7i, j**).

### Pharmacological inhibition of cGAS rescues neuronal transcriptomic changes induced by Dravet mutation

We next investigated the effects of cGAS inhibition on neuronal transcriptomics. In ENs, pseudobulk analysis revealed that cGASi treatment reversed the expression of 505 out of 992 differentially expressed genes (DEGs) (**Fig. 8a**, Table S6), with a strong negative correlation in logFC (*r* = -0.959) (**Fig. 8b**). Top DEGs included *Bdnf, Homer1, Satb2,* and *Mef2c* (**Fig. 8b**). Consistent with cGASi-mediated *Mef2c* upregulation^21^, enrichment analysis showed that 96 rescued DEGs were *Mef2c* targets (odds ratio 12.3), many of which regulate neuronal excitability, including neurotransmitter receptors (*Gria1, Grm1, Gria4*) and ion channels (*Camk2n1, Cacnb4, Kcnh5, Camk4, Kcnh7*, and others) (**Fig. 8c**). In INs, cGASi reversed the expression of 157 out of 476 DEGs (**Fig. 7d**, Table S6), with a similarly strong negative correlation (*r* = -0.956) (**Fig. 8e**). DEGs included genes involved in synaptic transmission (*Unc13c, Robo2*), and neuronal development, migration, and survival (*Runx2, Unc5c, Unc5d, Tenm3*) (**Fig. 8e**). Immunostaining confirmed the downregulation of MEF2C in *Scn1a+/-* mice, which was rescued by cGASi treatment (**Fig. 8f, g**).

**Figure. 8:**
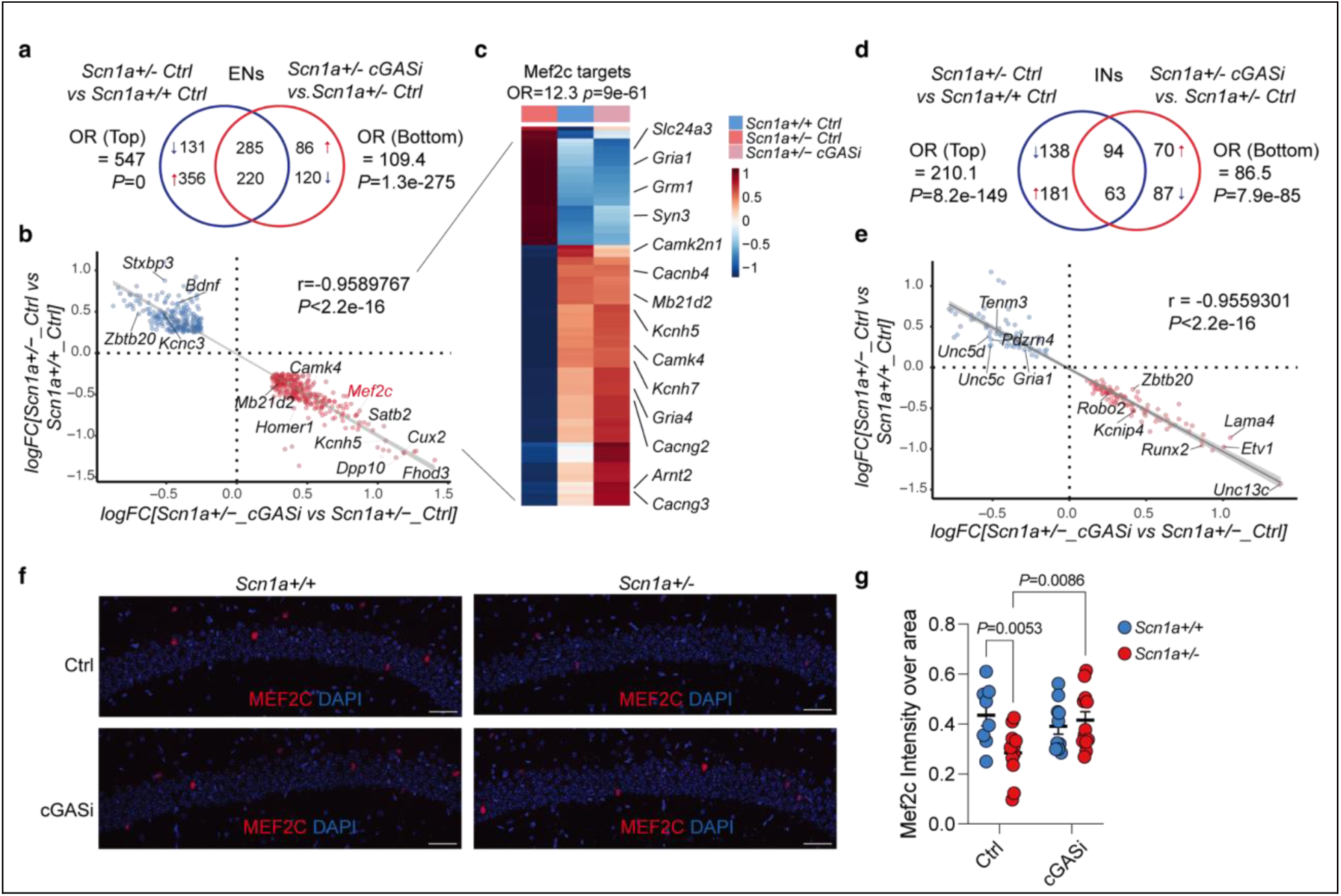
Pharmacological inhibition of cGAS rescues neuronal transcriptomic changes induced by Dravet mutation. (a) Venn diagram showing the overlap between the top DEGs (logFC >=0.25 or <= -0.25, *p*.adj <0.05) of *Scn1a+/-,* Ctrl versus *Scn1a+/+,* Ctrl and *Scn1a+/-*, cGASi versus *Scn1a +/-*, Ctrl excitatory neurons. Overlapping odds ratio and *P* value calculated with Fisher’s exact test. (b) Scatter plot and simple linear regression analysis with standard error showing a negative correlation between logFC values of overlapping DEGs in A). Pearson’s r = -0.96, *p*<2.2e-16. (c) Heatmap of the scaled average expression of *Mef2c* target genes in *Scn1a+/+,* Ctrl; *Scn1a+/+,* Ctrl; and *Scn1a+/-*, cGASi excitatory neurons. Overlapping odds ratio and *P* value showing the enrichment of *Mef2c* target genes in overlapping DEGs in B), Fisher’s exact test. (d) Venn diagram showing the overlap between the top DEGs (logFC >=0.25 or <= -0.25, *p*.adj <0.05) of *Scn1a+/-,* Ctrl versus *Scn1a+/+,* Ctrl; and *Scn1a+/-*, cGASi versus *Scn1a +/-*, Ctrl inhibitory neurons. Overlapping odds ratio and *P* value calculated with Fisher’s exact test. (e) Scatter plot and simple linear regression analysis with standard error showing a negative correlation between logFC values of overlapping DEGs in D). Pearson’s r = -0.96, *p*<2.2e-16. (f) Representative 40x immunofluorescence images of MEF2C in the CA1 region of hippocampi of N1 mice treated with cGASi or control. Scale bar, 50⎧m. (g) Quantification of MEF2C intensity over the CA1 area. Each circle represents the mean quantification of three to four brain sections per animal. Two-way ANOVA mixed-effects analysis with Fisher’s LSD test. *Scn1a+/+,* Ctrl, n=8; *Scn1a +/-*, Ctrl, n=11; *Scn1a+/+,* cGASi, n=10; *Scn1a+/-,* cGASi, n=12.

We then compared cGASi effects with genetic *Cgas* knockdown by analyzing overlaps in DEGs from N3 *Scn1a+/-, Cgas+/-* vs. *Scn1a+/-* ENs and N1 *Scn1a+/-, cGASi* vs. *Scn1a+/-, Ctrl* ENs. Despite differences in genetic background and mouse age, 147 common DEGs were identified, with a correlation of *r* = 0.486 (**Extended Data Fig. 12a**). Key shared DEGs included *Homer1, Pde10a, Satb2,* and *Bdnf*, highlighting their importance in epilepsy (**Extended Data Fig. 12b**). In summary, cGAS knockdown and inhibition exerted similar effects by correcting a major portion of the neuronal transcriptomic alterations induced by DS.

## DISCUSSION

Our study identifies IFN-I signaling as a key inflammatory pathway contributing to disease progression in drug-refractory epilepsy (DRE), particularly in Dravet syndrome (DS). Transcriptomic analysis of human DRE tissue and DS mouse models revealed elevated IFN-I signatures in microglia, highlighting sustained neuroinflammation as a hallmark of pharmacoresistance. This aligns with observations of persistent inflammation in patients with prolonged seizures and provides molecular insights into resistance to antiepileptic drugs (AEDs). Notably, IFN-I signaling mediates a unique inflammatory response in epilepsy, distinct from other disease-associated microglial (DAM) signatures, underscoring its specific role in epileptogenesis.

Our findings further implicate cGAS, a cytosolic DNA sensor, as a critical upstream regulator of IFN-I signaling in epilepsy. Activation of the cGAS-STING pathway was elevated in both human DRE tissue and DS mouse models, likely driven by neuronal DNA damage induced by neuronal activity and exacerbated in pathological conditions^45, 46^. Supporting this, our *in vitro* studies showed that conditioned media from hyperactive neurons activated the cGAS pathway in microglia, linking neuronal hyperactivity to microglial IFN-I signaling. Because the human specimens were obtained from surgical pathology and autopsy archives, the control and epilepsy groups were not age matched, and the precise cortical regions available for analysis varied across individuals. Future studies using larger, age and region matched human cohorts will be essential to rigorously quantify alterations in the cGAS STING pathway in drug resistant epilepsy.

Clinical evidence reinforces this connection: patients with type I interferonopathies like Aicardi-Goutières syndrome often develop epilepsy^47^, and IFN-I therapy for chronic viral hepatitis can induce seizures that resolve upon cessation of treatment^48–50^. Activation of interferon signaling is also identified in DRE associated with tuberous sclerosis complex^51^. IFN-I signaling is triggered by pattern-recognition receptors (PRRs) responding to DAMPs and PAMPs, with nucleic acid-sensing PRRs such as TLR3 and TLR9 previously implicated in seizure models^11, 16^. Our findings expand on this by positioning cGAS as a central mediator of neuroinflammation and a potential therapeutic target in epilepsy.

We found that targeting cGAS effectively reduced epilepsy phenotypes across multiple models. Genetic deletion of cGAS lowered the severity of and increased latency to seizures in pharmacologically induced PTZ and KA models and reduced hyperthermia-induced seizure susceptibility and spontaneous epileptiform activity in two DS mouse lines with different genetic backgrounds. Our findings indicate that recurrent seizures drive sustained activation of the cGAS STING pathway, which in turn contributes to the maintenance of the epileptic state in a chronic mouse model. In addition, aberrant activation of cGAS STING exacerbates seizure severity in acute PTZ and kainic acid induced models, highlighting its dual role as both a downstream consequence of seizure activity and an upstream amplifier of epileptogenic processes. cGAS reduction also diminished IFN-I signaling and pro-inflammatory pathways such as antigen presentation, complement activation, and TLR signaling in microglia, as well as disease-associated astrocyte (DAA) signatures. Astrocytes, which interact with microglia and neurons, promote epileptogenesis through neuroinflammation, lipid metabolism, and gliotransmission^25, 52, 53^. We observed that cGAS reduction decreased APOE accumulation in DS astrocytes, aligning with reports that APOE-high reactive astrocytes exacerbate neuronal hyperactivity and epilepsy progression^25^. Although widely used for screening anti-epileptic drugs, induced seizure models assess seizure susceptibility and severity under provoked conditions and do not fully recapitulate endogenous spontaneous seizure activity. Therefore, to better reflect the etiological and phenotypic diversity of complex human epilepsies, future studies employing complementary models will be necessary to determine whether this cGAS reduction effects extend to other seizure types.

In neurons, cGAS reduction restored transcriptomic changes related to synaptic transmission, ion channels, and neurotransmitter receptors but did not affect *Scn1a* expression or Nav1.1 protein levels, suggesting a mechanism not specific to DS. Neuronal DNA damage, mtDNA defects, and DNA release—common in epilepsy^15–17^—likely activate cGAS. Elevated 53BP1 levels in neurons from human DRE tissue confirmed increased DNA damage. *In vitro*, conditioned media from PTZ-treated neurons increased TBK1 phosphorylation in *Cgas+/+* but not *Cgas-/-* microglia, linking neuronal DNA release to cGAS activation. In contrast, direct PTZ exposure failed to induce pTBK1 phosphorylation in primary microglia, arguing against a cell autonomous effect of PTZ on cGAS STING activation. Consistent with this, our in vivo analyses in DS mice demonstrate increased microglial cytosolic dsDNA and extracellular vesicle associated DNA following seizure activity, providing direct evidence that hyperactive neurons indirectly drive microglial cGAS STING activation in vivo. This aligns with previous studies showing that self-DNA from degenerating neurons activates DNA sensors like TLR9 during seizures^16^. Additionally, IFN-I-responsive microglia specifically engulfing neurons with DNA damage further supports this mechanism^26^.

Our findings support that targeting the cGAS-STING-IFN-I pathway offers promise for treating DS. The brain-permeable cGAS inhibitor TDI-6570 reduced early mortality, hyperthermia-induced seizure susceptibility, and corrected many transcriptomic changes in DS mice. Microglial inflammatory signatures varied between young (N1) and older (N3) DS mouse lines, reflecting differences in disease stage and genetic background. Nonetheless, the consistent elevation of cGAS in DS models highlights its role in acute and chronic epilepsy. TDI-6570 also increased *Mef2c* expression, a transcription factor critical for synapse regulation and neuronal development^43, 54, 55^, and rescued the expression of its target genes. Given that MEF2C haploinsufficiency syndrome includes epilepsy as a common symptom^20^ and neuronal MEF2C is negatively regulated by STING-IFN-I activation^21^, the rescue of MEF2C transcriptional network likely contributes to the protective effects of cGAS inhibition. Pharmacological inhibition of cGAS was notably more effective than genetic reduction in improving survival and hyperthermia-induced seizure thresholds. This difference may stem from variations in timing, level, and mode of inhibition. Genetic reduction occurs embryonically, whereas cGASi is administered postnatally. IFN-I-responsive microglia is crucial in maintaining excitatory/inhibitory balance in neurodevelopment^26^, and this function may be disrupted by genetic reduction of cGAS. Further, compensatory changes may arise with genetic manipulation but not with inhibitor treatment. Additionally, *Cgas+/-* reduces both enzymatic and non-enzymatic functions of cGAS, while cGASi specifically targets enzymatic activity. The role of cGAS’s non-enzymatic functions remains less understood.

The exact cell type(s) in which cGAS activity drives disease progression in DS remains an important unanswered question. Microglia express the highest levels of cGAS and STING and play a central role in DNA-sensing and neuroimmune signaling^21^. Microglia cGAS-STING activation triggers IFN-I production, which disseminates antiviral responses to other CNS cell types. For instance, during HSV-1 infection, microglial cGAS-STING activation induces IFN-I production, which activates STING-dependent antiviral programs in neurons and primes TLR3-IFN-I pathways in astrocytes^56^. Similarly, STING-mediated inflammatory responses have been shown to propagate in CNS and peripheral neurons^57–59^. Our snRNAseq data supports these findings, showing that *Tlr3* is upregulated in *Scn1a+/-* astrocytes, a change reversed by *Cgas+/-*. Furthermore, a microglial-specific Cre-induced cGAS activation model demonstrated heightened IFN-I responses in both microglia and astrocytes, which was sufficient to cause cognitive impairments^60^. We further observed that PTZ-induced seizure activity was reduced in mice with microglia-specific cGAS deletion, supporting a role for microglia cGAS STING signaling in seizure expressions. In DRE, it is conceivable that microglial cGAS activation by engulfed neuronal DNA drives IFN-I production, amplifying inflammatory responses in neurons and astrocytes. This exacerbates neuronal hyperexcitability, leading to neuronal damage and creating a feed-forward cycle of inflammation and sustained hyperexcitability. Suppressing cGAS activation caused by dsDNA released from hyperexcitable neurons could disrupt this vicious cycle, potentially halting DRE progression. Future studies employing conditional knockout models and systematically investigating the cell-type-specific roles of cGAS in the CNS are essential to fully elucidate its contributions to epilepsy. These efforts could advance our understanding of cGAS as a therapeutic target and clarify its broader role in neuroinflammatory and neurological diseases.

## METHODS

### Mice

Mice were housed no more than five per cage, given ad libitum access to food and water and housed in a pathogen-free barrier facility at 21–23 °C with 30–70% humidity on a 12-h light/12-h dark cycle. All mouse protocols were approved by the Institutional Animal Care and Use Committee at Weill Cornell Medicine.

For the conditional DS snRNAseq, frozen whole brains of *PV-cre/+, Scn1a+/+* (n=3) and *PV-cre/+, Scn1a F/+* (n=3) mice on the C57BL/6J background were obtained from Dr. Jorge Palop (University of California, San Francisco).

For the pharmacological model of seizure, *Cgas−/−* mice (The Jackson Laboratory, 026554) were crossed with C57BL/6J mice (The Jackson Laboratory, 000664) to generate F1 *Cgas+/−* mice. F1 *Cgas+/−* mice were then crossed again to generate littermate *Cgas+/+*, *Cgas+/-*, and *Cgas-/-*mice. For targeting microglia-specific cGAS deletion in adult mice, *Cx3cr1^CreER^* mice^61^ (The Jackson Laboratory, 020940) were crossed with a floxed *cGas^tm^*^1c^ mice^62^ (MMRRC, 065342-UCD) to generate *cGas^tm^*^1c^*: F/F; CX3CR1^CreER^ Cre/+* and littermate *cGas^tm^*^1c^ *F/F; CX3CR1^CreER^ +/+* control mice. Both groups received tamoxifen injection (20mg/kg, i.p.) once daily for 5 consecutive days at 2 months of age.

For the DS model, female *Cgas+/−* mice were crossed with male *129S-Scn1a^tm1Kea^ /Mmjax* (The Jackson Laboratory, 037107) to generate N1 *Scn1a+/+*, *Scn1a+/-*, *Scn1a+/+, Cgas+/−*, and *Scn1a+/-, Cgas+/-* mice. Mice of both sexes were included for mortality, hyperthermia-induced seizure, histological and biochemical analyses.

A separate breeding of female *Cgas−/−* mice with male *129S-Scn1a^tm1Kea^ /Mmjax* were used to generate N1 *Scn1a+/-, Cgas+/-* mice, which were crossed again with *Cgas−/−* mice to generate N2 *Scn1a+/-, Cgas+/-* and *Scn1a+/-, Cgas-/-* mice. Male N2 *Scn1a+/-, Cgas+/-* mice were crossed with female C57BL/6J mice to generate N3 *Scn1a+/+*, *Scn1a+/-*, *Scn1a*+/+, *Cgas+/−*, and *Scn1a+/-, Cgas+/-* mice. Mice of both sexes were included in analysis. Mice that received electrode implantation surgeries were used for LFP recording and subsequent hyperthermia-induced seizure induction and were excluded from mortality, transcriptomic, histologic, and biochemical analysis.

For cGASi treatment, male *129S-Scn1a^tm1Kea^ /Mmjax* mice were crossed with female C57BL/6J mice to generate N1 *Scn1a+/+* and *Scn1a+/-* mice. Mice of both sexes were included for mortality, hyperthermia-induced seizure, transcriptomic, histological and biochemical analyses.

### Nuclei isolation from frozen mouse brain

Nuclei isolation from frozen mouse brain was adapted from a previous study, with modifications^63, 64^. All procedures were performed on ice or at 4 °C. In brief, postmortem brain tissue was placed in 1,500 µl of Sigma nuclei PURE lysis buffer (Sigma, NUC201-1KT) and homogenized with a Dounce tissue grinder (Sigma, D8938-1SET) with 20 strokes with pestle A and 15 strokes with pestle B. The homogenized tissue was filtered through a 35-µm cell strainer, centrifuged at 600g for 5 min at 4 °C and washed three times with 1 ml of PBS containing 1% bovine serum albumin (BSA), 20 mM DTT and 0.2 U µl−1 recombinant RNase inhibitor. Nuclei were then centrifuged at 600g for 5 min at 4 °C and resuspended in 800 µl of PBS containing 0.04% BSA and 1× DAPI, followed by fluorescence-activated cell sorting to remove cell debris. The sorted suspension of DAPI-stained nuclei was counted and diluted to a concentration of 1,000 nuclei per µl in PBS containing 0.04% BSA.

### Droplet-based snRNA-seq

For droplet-based snRNA-seq, libraries were prepared with Chromium Single Cell 3′ Reagent kits (v3.1; 10x Genomics, PN-1000268), according to the manufacturer’s protocol. The snRNA-seq libraries were sequenced on a NovaSeq 6000 sequencer (Illumina) with PE 2 x 50 paired-end kits by using the following read length: 28 cycles Read 1, 10 cycles i7 index, 10 cycles i5 index, and 90 cycles Read 2.

### Analysis of droplet-based snRNA-seq data

In 6-month-old conditional Dravet mice, we sequenced and integrated samples from the frontal cortical and hippocampal tissue of *PV-cre/+, Scn1a +/+* (n = 3) and *PV-cre/+, Scn1a F/+* (n = 3) mice. In 4-5-month-old N3 Dravet cohort, we sequenced and integrated samples from the hippocampi of *Scn1a +/+* (n = 4), *Scn1a +/-* (n = 4), *Scn1a+/+, Cgas+/-* (n = 4), and *Scn1a+/-, Cgas+/-* (n=4) mice. In 1.5-month-old N1 Dravet cGASi treatment cohort, we sequenced and integrated samples from the hippocampi of *Scn1a +/+,* Ctrl (n = 4), *Scn1a +/-*, Ctrl (n = 4), *Scn1a+/+,* cGASi (n = 4), and *Scn1a+/-,* cGASi (n=4) mice. Gene counts were obtained by aligning reads to the mm10 genome with Cell Ranger software (v6.1.2; 10x Genomics). To account for unspliced nuclear transcripts, reads mapping to pre-mRNA were counted. Cell Ranger 6.1.2 default parameters were used to call cell barcodes. We further removed genes expressed in no more than 3 cells, cells with unique gene counts over 5,000 or less than 300, cells with UMI counts over 15,000 and cells with a high fraction of mitochondrial reads (>1% or >5% for the N3 cohort). Potential doublet cells were predicted using DoubletFinder^65^ for each sample separately, with high-confidence doublets removed. Normalization and clustering were done with the Seurat package v5.0.0. In brief, counts for all nuclei were scaled by the total library size multiplied by a scale factor (10,000) and transformed to log space. A set of 2,000 highly variable genes was identified with SCTransform from the SCTransform R package in the variable stabilization mode. This returned a corrected unique molecular identifier count matrix, a log-transformed data matrix and Pearson residuals from the regularized negative binomial regression model. Principal-component analysis was done on all genes, and t-distributed stochastic neighbor embedding was run on the top 15 principal components.

The major cell types in the brain were identified by the expression of known cell identity markers^10, 66, 67^: excitatory neurons (*Slc17a7, Nrgn*), inhibitory neurons (*Gad1, Gad2*), immune cells (*Cx3cr1*, *P2ry12*, and *Csf1r*), oligodendrocytes (*Plp1*, *Mbp*, and *Mobp*), oligodendrocyte precursor cells (*Vcan*, *Pdgfra*), astrocytes (*Clu*, *Plpp3*, and *Pla2g7*), vascular cells (*Bmp6*, *Adam12*, *Cped1*), and choroid plexus epithelial cells (*Clic6*, *Ttr*). Cell clusters were identified with the Seurat functions FindNeighbors and FindClusters. In this analysis, the neighborhood size parameter pK was estimated using the mean variance-normalized bimodality coefficient (BCmvn) approach, with 15 principal components used and pN set as 0.25 by default. For each cluster, we assigned a cell-type label using statistical enrichment for sets of marker genes and manual evaluation of gene expression for small sets of known marker genes^10, 67–69^.

Differential gene expression analysis was done using the FindMarkers function and MAST^70^. To identify gene ontology and pathways enriched in the DEGs, DEGs were analyzed using the clusterProfiler package in R (v4.10.1) or the MSigDB gene annotation database^71, 72^. To identify gene activation networks and upstream regulators, DEGs were analyzed using Ingenuity Pathway Analysis (QIAGEN, Inc.). To control for multiple testing, we used the Benjamini–Hochberg approach to constrain the FDR. The overlapping odds ratio and *P* value between DEG datasets were calculated with GeneOverlap package in R (v1.38.0).

SCENIC^27^ (Single Cell rEgulatory Network Inference and Clustering) analysis was performed in R to interrogate gene regulatory networks/transcription factor activity in the conditional Dravet microglia clusters. Analysis was performed following the tutorial on https://github.com/aertslab/SCENIC.

hdWGCNA^28^ analysis was performed in R to identify gene module networks in the microglia clusters of conditional Dravet and N3 Dravet datasets. Analysis was performed following the tutorial on https://smorabit.github.io/hdWGCNA/articles/basic_tutorial.html. For the conditional Dravet microglia cluster, “Seurat clusters” was used to construct the metacells. For the N3 Dravet microglia cluster, “genotype” was used to construct the metacells. Differential module eigengene (DME) analysis was performed to identify modules that are up-or down-regulated in specified groups of cells. The EnrichR package was further used to identify pathways highly expressed in each module.

### Immunofluorescence

Hemibrains from transcardially perfused mice were placed in 4% paraformaldehyde for 28 h, followed by 30% sucrose in PBS for 48 h at 4 °C. Sections were cut coronally at 40 µm using a freezing microtome (Leica) to produce eight to ten free-floating series per mouse and placed in cryoprotective medium at –20 °C until use. All washing steps were 5 min long and repeated three times. Sections were washed in 1xTris-Buffered Saline with Tween 20(TBST), permeabilized with 0.5% Triton X-100 in TBST for 10 min and washed. Sections were then placed in an antigen-retrieval solution (Antigen Decloaker, Biocare Medical; CB910M) and placed in a 90 °C incubator for 15 min, when applicable. Sections were washed and then blocked in 5% normal donkey serum (Vector, BMK-2202) in TBST for 2 h at room temperature. Primary antibodies were diluted in 5% normal donkey serum in TBST and incubated overnight at 4 °C. The following day, sections were washed thoroughly and incubated in appropriate secondary antibodies (1:500; Invitrogen) for 1 h. Sections were washed, mounted on slides using Vectashield antifade mounting medium with DAPI (Vector Laboratories, H-1200; or Sigma, D9542), and imaged.

5 µm paraffin-embedded human brain sections mounted on slides were incubated in oven at 60°C for 10 min with aluminum foil underneath to catch melted paraffin. Slides were deparaffinized with Xylene for 5 mins for three times followed by 100% EtOH for 2 mins for 2 times, 95% EtOH for 2-5 mins, and washed three times in dH_2_O for 2 mins each. Slides were then placed in an antigen-retrieval solution made of 1x Citrate Buffer, pH6.0 (Electron Microscopy Sciences, Cat # 64142-08) in dH_2_O and placed in a pressure cooker (Cuisinart) at high pressure for 15 mins. Slides were washed for three times of 2min each with cool dH_2_O followed by three washes of 2min each with 1x PBS. Slides were then blocked with 5% donkey serum in 1x PBS for 1 hr and incubated in primary antibodies diluted in 1% donkey serum in 1x PBS overnight in a humidified slide chamber. The following day, slides were washed in 1x PBS for 2 mins for three times and incubated with secondary antibodies diluted in 1% donkey serum in 1x PBS for 1 hour in a humidified slide chamber. Slides were then washed in 1x PBS for 2 mins for three times and coverslipped with Vectashield mounting medium with DAPI.

Primary antibodies used included anti-STING (clone D2P2F, Cell Signaling Technology, 13647; 1:200), anti-IBA1 (Abcam, ab5076; 1:300; or Wako, 019-19741; 1:1000), anti-NeuN (clone A60, Millipore, MAB377; 1:500), anti-GFAP (Abcam, ab7260; 1:1000), anti-APOE (a generous gift from Dr. Holtzman (Washington University in St. Louis); 1:300), anti-PDE10A (Abcam, ab227829; 1:500), anti-MEF2C (clone 681824, R&D Systems, MAB6786; 1:200), anti-53BP1 (Novus Biological, NB100-304; 1:1000), anti-MAP2 (Novus Biological, NB300-213; 1:1000); anti-dsDNA (Abcam, ab27156; 1:2000). Secondary antibodies used were Alexa Fluor donkey anti-rabbit/goat/chicken 488, anti-mouse 568 and donkey anti-goat 568 (Invitrogen) at a dilution of 1:500, or Alexa Fluor goat anti-rabbit 488 and anti-mouse 594 (Invitrogen) at a dilution of 1:800.

For all mouse brain staining, 3-4 sections per animal were imaged with a Zeiss Apotome ×20 objective (Carl Zeiss). For IBA1, STING; GFAP, APOE; and PDE10A, NeuN staining, the CA1 or CA3 region of the hippocampus was imaged with a 1-μm interval z-stack over a total distance of 8 μm per slice. For MEF2C staining, the CA1 region was imaged with 4×1 tiles and a 1-μm interval z-stack over a total distance of 10 μm per slice. Final images were processed with maximum intensity projection. For all human brain imaging, 4-5 images over different areas per section were acquired with a Keyence BZ-X700 microscope using a ×20 objective. Quantification was done with ImageJ software V2.1.40LL (NIH).

All images of dsDNA staining were acquired using a Nikon A1 Laser Confocal microscope. The large images containing hippocampus were obtained by tile-scan under 20x magnification. Single z-stack areas of hippocampus CA1 were imaged under 100x magnification. 6 windows inside CA1 area were randomly selected for z-stack imaging under identical conditions. Images were processed and analyzed in Fiji-ImageJ. For dsDNA puncta in MG cytoplasm, z-stack images of 10 layers with 1 um z-step containing a single microglia cell were projected by maximum intensity. dsDNA puncta inside the MG cell body and main branches but outside its nucleus were quantified as “cytosolic dsDNA puncta number” in single microglia.

### Tissue EV Isolation

The tissue samples were digested by collagenase, Type 3 (Cell Signaling Technology, 25970S) in Hibernate-E medium (Thermo Fisher A12476-01) at 37 ^0^C for 15 minutes. The enzymatic reaction was stopped by adding cOmplete protease inhibitor (Millipore Sigma. 11697498001) and PhosSTOP phosphatase inhibitor (Millipore Sigma, 4906837001). The dissociated tissue was centrifuged at 300g for 10 min, 2000g for 15 min, and 10,000g for 30 min at 4 °C with the supernatant from each step. Supernatants from the 10, 000g step were subjected to EV isolation using Izon size exclusion columns (qEV Original, 70 nm). EV-enriched fractions were ultracentrifuged at 100,000 × g at 4 °C for 70 minutes, and the resulting pellets were resuspended in DPBS^73^.

### Nano-flow cytometry measurement (NanoFCM)

EV samples were analyzed by nano flow cytometry to quantify particle concentration and size distribution. Tissue EV samples were diluted 1:50 in HEPES buffer. Instrument calibration for particle concentration was performed using QC beads, and size calibration was performed using a silica nanosphere cocktail (S16M-Exo). Events were recorded during a 1 min acquisition period for each sample. Particle concentration and size were calculated from flow rate and side-scatter intensity using NanoFCM Profession software (v3.0)^73, 74^.

### Western blots

For mouse brain samples, half hippocampi were sonicated on ice in RIPA buffer (Thermo Fisher Scientific, 89901) supplemented with Halt phosphatase inhibitor cocktail (Thermo Fisher Scientific, 78420) and cOmplete protease inhibitor cocktail (Roche, 11697498001). After sonication, brain lysates were centrifuged at 20,000 g for 15 min at 4°C. Supernatants were collected, and protein concentration was measured by BCA (Pierce, 23225). Samples were boiled at 95 °C for 5 min or 55 °C for 30 min (for Nav1.1 only). 50 micrograms of protein were loaded onto NuPage Bis-Tris gels (Thermo Fisher) and run in SDS running buffer at 80 V for 30 minutes and 130 V for ∼1 hour. Gels were transferred to PVDF membranes (Bio-Rad) for 1.5h at 0.36A in a cold room. Membranes were blocked for 1 h in 5% milk in TBST. Primary antibodies were diluted in 1% milk in TBST and incubated at 4 °C overnight. The following day, membranes were washed three times for 5 min each in TBST and incubated with secondary antibodies in 1% milk in TBST for 1 h at room temperature. Membranes were washed three times, then treated with ECL (Bio-Rad) for 60 s and developed using a Bio-Rad imager. Blots were scanned at 300 d.p.i. and quantified using Image Lab.

Primary antibodies used for western blotting were cGAS (clone D3O8O, Cell Signaling Technology, 31659; 1:500), STING (clone D1V5L, Cell Signaling Technology, 50494; 1:1000), TBK1 (clone D1B4, Cell Signaling Technology, 3504; 1:1,000), pTBK1 (clone D52C2, Cell Signaling Technology, 5483; 1:500), GAPDH (GeneTex, GTX100118; 1:10,000), Nav1.1 (clone K74/71, NeuroMab; 1:500), GFAP (Abcam, ab7260; 1:1000), β-Actin (Millipore Sigma, A3854, 1:10,000) . Secondaries used for western blotting were Peroxidase-AffiniPure Goat anti-Rabbit IgG (Jackson ImmunoResearch Labs, 111-035-144; 1:10,000) or Peroxidase-AffiniPure Goat anti-Mouse IgG (Jackson ImmunoResearch Labs, 115-035-146; 1:10,000).

### Pharmacological model of seizures

Pentylenetetrazol (PTZ) powder (Sigma-Aldrich, P6500) and kainic acid (KA) monohydrate (Tocris, 7065) was dissolved in 0.9% sodium chloride injection, USP (Hospira, Inc) to a final concentration of 5mg/ml (PTZ) and 2.5 mg/ml (KA), respectively. Mice were placed in a circular open field chamber (8228, Pinnacle Technology, Lawrence, KS) and video recorded with a FLIR camera (Teledyne FLIR, North Billerica, MA) for baseline activity for 10 minutes. Mice were then injected intraperitoneally with the PTZ solution at 50mg/kg or KA solution at 18mg/kg and observed simultaneously by 2 investigators under video for 30 minutes (PTZ) or 90 minutes (KA) to assess seizure activity. Seizures were scored based on a modified Racine scale^32^, where 0: no seizure response, 1: sudden behavioral arrest or immobility, 2: tail twitching, myoclonic jerks, or facial automations, 3: head nodding and neck jerks, 4: clonic seizures while the animal falls into a sitting position, 5: tonic-clonic seizures with the animal falling on its side, 6:wild-jumping, 7: tonic extension of limbs, 8:death. Maximum Racine score and latency to seizure were analyzed.

### Survival monitoring

Mice were monitored daily for survival starting from P20 and individual animal status was recorded using Mouse Laboratory Information System (mLIMS). Survival data were analyzed for each genotype and survival curves were generated using Prism (GraphPad Software, Inc., CA, USA).

### Hyperthermia-induced seizure

Hyperthermia-induced seizure was performed based on a previously published protocol with modifications^75^. Mice were placed in an open field chamber with a RET-4 probe (Phisitemp) inserted in the rectum to continuously measure rectal temperature. Mice were allowed to habituate and recorded at baseline for 10 min, then heated with an infrared heat lamp (HL-1, Phisitemp) at a constant distance from the chamber to gradually increase the body temperature using a TCAT-2DF thermo controller (Phisitemp). For every 0.5 °C the body temperature was increased, the heat lamp was turned off for 30s. The cycle repeated until clonic or tonic-clonic seizure was recognized by LFP recording (in N3 mice) and video analyses (in both N1 and N3 mice) or the body temperature reached 42.5 °C. The heating lamp was then switched off and an ice block was placed in the chamber to allow recovery. Mice were monitored until the body temperature returned to the baseline or until death occurred. For tissue EV and dsDNA immunofluorescence analysis, brains were collected 3 hours following hyperthermia-induced seizures (**Fig. 3a**).

### 24-hour local field potential recording in freely moving mice

Mice were initially anesthetized with 3.5 % isoflurane mounted on a stereotaxic frame and maintained under ∼1.2% isoflurane. Body temperature was maintained at 37 °C with a regulated heating blanket (World Precision Instruments, Sarasota, FL). A craniotomy was drilled for electrode insertion. Animals were implanted with a custom 32-ch 3D electrode array (Kedou Brain-Computer Technology, Suzhou, China) that contains 8 tetrodes over somatosensory (S1), the CA1 of hippocampus, dentate gyrus (DG), primary visual cortex (V1) and thalamus (TH) (S1 : antero-posterior (AP) -1.8mm, mediolateral (ML) 2.1 and 2.3mm, dorsoventral (DV) 1.0mm, CA1: AP -1.8mm, ML 1.5mm, DV1.5mm, DG: AP -3.0mm, ML 1.8mm, DV 2.2mm, V1: AP -3.0mm, ML 2.1 and 2.3mm, DV 0.8mm. TH: AP -1.8mm, ML 1.0 and 1.2 mm, DV 3.8 mm. Fig. 2K). Stainless steel screws were implanted into the skull to provide electrical ground and mechanical stability for drives and the whole construct was bonded to the skull using C&B-Metabond luting cement (Parkell, Edgewood, NY). Two weeks after implantation, animals were put in a circular open field chamber (8228, Pinnacle Technology, Lawrence, KS) to record spontaneous activity continuously for 24 hours with food and water supply. Electrophysiological data were acquired using an Intan RHD headstage (Intan Technologies, LA, CA) and the Open Ephys acquisition board and software (OEPS, Alges, Portugal) sampled at 1kHz^76^. Locomotion was simultaneously acquired with a FLIR camera (Teledyne FLIR, North Billerica, MA) and the open-source Bonsia software^77^ recording at 50 Hz.

### Epileptiform activity analysis

The detection and quantification of epileptic activities were performed using a custom script written in Matlab based on an automated algorithm^78^(https://github.com/Jackson-Kyle-CCOM/Automated-EEG-Algorithm). LFP recording signals were firstly band-pass filtered between 5 and 90 Hz. The baseline activity value was autonomously calculated based on a period of 10 minutes of low activity. The lower threshold value was defined as 5 standard deviations from baseline value. An upper threshold value of 6000 μV was used to remove EEG artifacts due to noise and animal behavior. Then LFP spikes for whole recording were determined following those key parameters. The inter-spike interval was set at 100 ms and the maximum peak width was set to 200 ms to identify interictal spike, polyspikes, and other abnormal spiking patterns or LFP artifacts. Seizure-like ictal ^79, 80^ epileptiform was defined to contain sequential peaks that lasted for at least 10s with an inter-spike interval of less than 5s.

### Primary neuron isolation and culture

Primary neuron cells were isolated from mouse embryos at prenatal E16. Briefly, the brains were isolated and minced. Tissues were dissociated in 0.25% Trypsin-EDTA (Gibco, A5200-056) for 15 min at 37°C. For every 4 brains, 300uL DNase I (Millipore Sigma, 260913) was added to the tissue and the supernatant was discarded. Next, the tissue was washed twice with 2.7ml of HBSS (Gibco, 14025) and incubated for 5 mins at RT, the supernatant was discarded. 10ml of MEM (Gibco, 11095-080) with 10% FBS (Gibco, 26140-079) was then added and moderate pipetting was performed to dissociate the cells.

Primary neuron cells were then counted and seeded on 6 well plates with FBS-MEM, 1x B27 plus (Lifetechnology, A35828-01), and 1x N2 (Lifetechnology, 17502-048). Cells were maintained by half changing the media with 10μM Uridine and 5-FDU every 3-4 days.

### Primary neuron treatment with PTZ

On day 12 of the neuronal culture, 1.38g of PTZ powder (Sigma-Aldrich, P6500) was dissolved in 1ml DPBS to make a 10M PTZ stock solution. The cells were treated with 10mM PTZ in media for 1 hour, and then the media was removed, and cells were gently washed twice with fresh media. 3ml total of fresh and untreated maintenance media at a 1:1 ratio was then added to each well and collected after 24hrs of incubation as the neuronal conditioned media.

### Primary microglia isolation and culture

Brains were dissected from mouse pups at postnatal day 1-3. The meninges were removed, and the cortical tissue was finely chopped with a razor blade and digested in 2.5% trypsin (Gibco, 15090-45) with DNase I (Millipore Sigma, 260913) at 37 C for 22 minutes. The reaction was stopped with DMEM (Corning, 10-017-CV) containing 20% heat-inactivated FBS (Gibco, 26140-079). 200ul of DNAse was added followed by 5ml of 20% FBS/DMEM, and tissue was triturated and spun at 200 g for 15 minutes. The pellet was resuspended in DMEM supplemented with 10% heat-inactivated FBS (Gibco, 26140-079) and 1% penicillin-streptomycin (Gibco, 15070063) and was plated in T75 flasks pre-coated with 5 ug/ml poly-D-lysine (Thermo Fisher Scientific, A389401) and rinsed with water. Mixed glial cultures were maintained in growth media (DMEM with 10% FBS and 1% P/S) at 37 C and 5% CO2. On day 4, media was collected and spun down at 200rcf for 10mins to remove the debris. The supernatant was collected back to the flasks with fresh media. On day 7, media was supplemented with 5 ng/ml recombinant mouse GM-CSF (R&D Systems, 415-ML-020-CF) to promote microglial proliferation. When flasks were ready for harvest on day 10-13, to isolate microglia, the flasks were shaken at 400 rpm for 2 hours, media was spun down at 200 g for 15 minutes, and cells were plated for experiments on poly-D-lysine coated plates.

### Primary microglia treatment with HT-DNA and neuronal conditioned media

300-400k of *Cgas+/+* and *Cgas-/-* primary microglia cells were seeded on each well of a 6-well plate. For HT-DNA treatment, 2ug/ml of HT-DNA was transfected with lipofectamine 3000 (Thermo Fisher Scientific, L3000) in maintenance media. For the neuronal conditioned media treatment, 50% of the neuronal PTZ conditioned media and 50% of maintenance media were added along with lipofectamine 3000. After 16 hours of treatment, the microglia cells were collected and lysed with RIPA buffer supplemented with Halt phosphatase inhibitor cocktail (Thermo Fisher Scientific, 78420) and cOmplete protease inhibitor cocktail (Roche, 11697498001) for western blot analysis.

### Calcium imaging in primary neuron acute treatment with PTZ

For neuronal Ca2+ image experiments, neurons were transduced with hSyn-jGCaMP8f lentivirus ^81, 82^ at DIV 7. At DIV 14 primary neuron coverslips was placed into a glass-bottom chamber at 33 °C containing 10 mM PTZ Ca2+ imaging buffer (20 mM HEPES, 119 mM NaCl, 5 mM KCl, 2 mM MgCl2, 30 mM glucose, 2 mM CaCl2, 10 MH PTZ, pH 7.2–7.4). Fluorescence time-lapse images were collected on a Nikon FN1 microscope using a 60x, 1.0 NA objective (CFI APO 60XW NIR, Nikon) and a C-FL GFP filter cube. An X-CITE LED illuminator (Nikon) was used for excitation. Images were collected using an ORCA-Fusion CMOS camera (Hamamatsu) with 4 × 4 binning (576×576 pixel resolution, 16-bit grayscale depth, 0.43 m/pix) and NIS-Elements AR software (Nikon). Exposure time was set to 20 ms. For spontaneous activity, images were acquired at 30 Hz for 4 min. We selected ROIs covering all identifiable cell bodies using a semi-automated algorithm in NIS-Elements AR (Nikon). Further quantifications were performed using custom-written MATLAB (Mathworks) scripts. The fluorescence time course (Δ F/ F_0_) each image frame was calculated as (F_t_-F_0_)/ F_0_, where F_0_ was an average of all frames acquired during the first 0.5-s window as a baseline and F_t_ is the fluorescence intensity at a given time. Photobleaching was corrected by fitting exponentially weighted moving averages^83^.

### Primary microglia treatment with PTZ

On day 12 of the microglia culture, 1.38g of PTZ powder (Sigma-Aldrich, P6500) was dissolved in 1ml DPBS to make a 10M PTZ stock solution. The cells were treated with 10mM PTZ in media for 1 hour. After treatment, the media was removed, and cells were gently washed twice with fresh media. Subsequently, 3ml total of fresh and untreated maintenance media at a 1:1 ratio was then added to each well. Microglia cells were collected 24 hours later and lysed with RIPA buffer supplemented with Halt phosphatase inhibitor cocktail (Thermo Fisher Scientific, 78420) and cOmplete protease inhibitor cocktail (Roche, 11697498001) for western blot analysis.

### Synthesis of cGAS inhibitors

TDI-6570 was prepared as described previously^84^. A minor modification was made to the method to obtain intermediates of TDI -6570 in multigram quantities from the reaction mixtures; all intermediates were obtained by filtration using water and cold methanol. Purity of the products was confirmed by performing liquid chromatography-mass spectrometry analysis.

### Almond butter Formulation for cGASi (prepared fresh every day)

TDI-6570 powder (1 mg) was added to almond butter (5 g) (Barney Butter Almond Butter) and the mixture was thoroughly blended using a spatula in a glass beaker. The resulting TDI-6570-almond butter product was transferred to plastic cups at 0.5 g per 20g of mouse weight and placed in cages housing a maximum of 5 mice and is equivalent to 0.02 mg TDI-6570 **/** 0.1 g Almond butter or 5 mg/kg mouse weight.

### Intraperitoneal Injection Formulation for cGASi

TDI-6570 (1 mg) was dissolved in DMSO (60 µl) and diluted with PEG 400 (1.34 ml) to obtain a homogenous solution. Water (0.6 ml) was added in portions to the above solution with a thorough mixing by vortexing. The resulting clear solution of TDI-6570 was administered to mice at 10ul per gram of mouse weight, which is equivalent to 5 mg/kg mouse weight.

### Multiplex ELISA with brain lysates

300 ug of proteins from hippocampal tissue previously lysed for western blots were evenly loaded based on protein concentration and measured using a customized ProcartaPlex Mouse Chemokine Panel multiplex kit (Thermo Fisher Scientific) according to the manufacture’s manual and read with the Luminex™ xMAP™ INTELLIFLEX System. Final cytokine concentrations were calculated with the MILLIPLEX Analyst software.

### Statistical Analysis

All data are reported as mean ± S.E.M. Statistical analyses were performed using R v4.3.3 and GraphPad Prism v.10.2.0. For data analysis using GraphPad, significant outliers were removed using Prism’s outlier calculator with ROUT test (Q=0.1%) and Grubb’s test (alpha=0.05). Shapiro-Wilk and Kolmogorov-Smirnov tests were performed to determine the normality of all data. Normally distributed data was analyzed using ANOVA or t-test. Non-normal data was analyzed using Kruskal-Wallis or Mann-Whitney tests. Morality and hyperthermia-induced seizure data were analyzed with Log-rank Mantel-Cox test. Gene set overlap analysis was performed with Fisher’s exact test using the GeneOverlap package in R^39^. All statistical details can be found in the figures and figure legends. A subset of mice from the immunohistochemistry cohort was randomly selected for snRNA-Seq studies. Animals were blinded from genotypes when performing image analysis and quantification for all immunohistochemistry experiments.

## DATA AVAILABILITY

All data associated with this study are in the paper or the Supplementary materials. All RNA-seq data will be deposited to the Gene Expression Omnibus (GEO) and accession codes will be available before publication.

## CODE AVAILABILITY

All custom codes used for snRNA-seq data analysis will be available at https://github.com/lifan36 before publication.

## ACKNOWLEDGEMENTS

This work was supported by the NIH (R01AG076448, R01AG072758, R01AG079557-01, and 1R01AG079291-01A1, to L.G.; R01AG074541 to L.G. and S.C.S; K99AG078493 to S.A.; R01NS145443 to J.W. and L.G.), the Rainwater Charitable Foundation (to. L.G) and Freedom Together Foundation (to L.G.), Cure Alzheimer’s Fund (to L.G. and S.C.S.), Daedalus Fund (to L.G. and S.C.S.), BrightFocus Foundation Postdoctoral Fellowship in Alzheimer’s Disease Research (A20201312F to S.A.), JumpStart Research Career Development Program (to S.A.).

## AUTHOR CONTRIBUTIONS

L.G. and Y.H. conceived the project and planned the experiments. L.G., Y.H., M.Z., J.W., M.Y.W., M.C., and S.C.S. designed experiments. Y.H., L.F., M.Y.W., Z.L., B.K., D.Z., M.C., R.K.N., K.N., P.Y., Y. J., H.L., J. Z., M.B., S.A., and M.Z. performed experiments or analyses. Y.H., M.Z., L.F., M.Y.W., M.C., R.K.N., S.A., S.C.S, H.C, W.J., S.G. developed experimental protocols, tools or reagents. Y.H., P.Y., and M.B. maintained the mouse colony. B.L.L. provided human brain sections. K.L. and J.J.P. provided conditional Dravet mouse brains. Y.H. and L.G. wrote the manuscript. All authors read and approved the manuscript.

