## Supplemental Data for "cGAS-mediated IFN-I signaling contributes to disease progression in drug-refractory epilepsy"

### EXTENDED DATA FIGURES

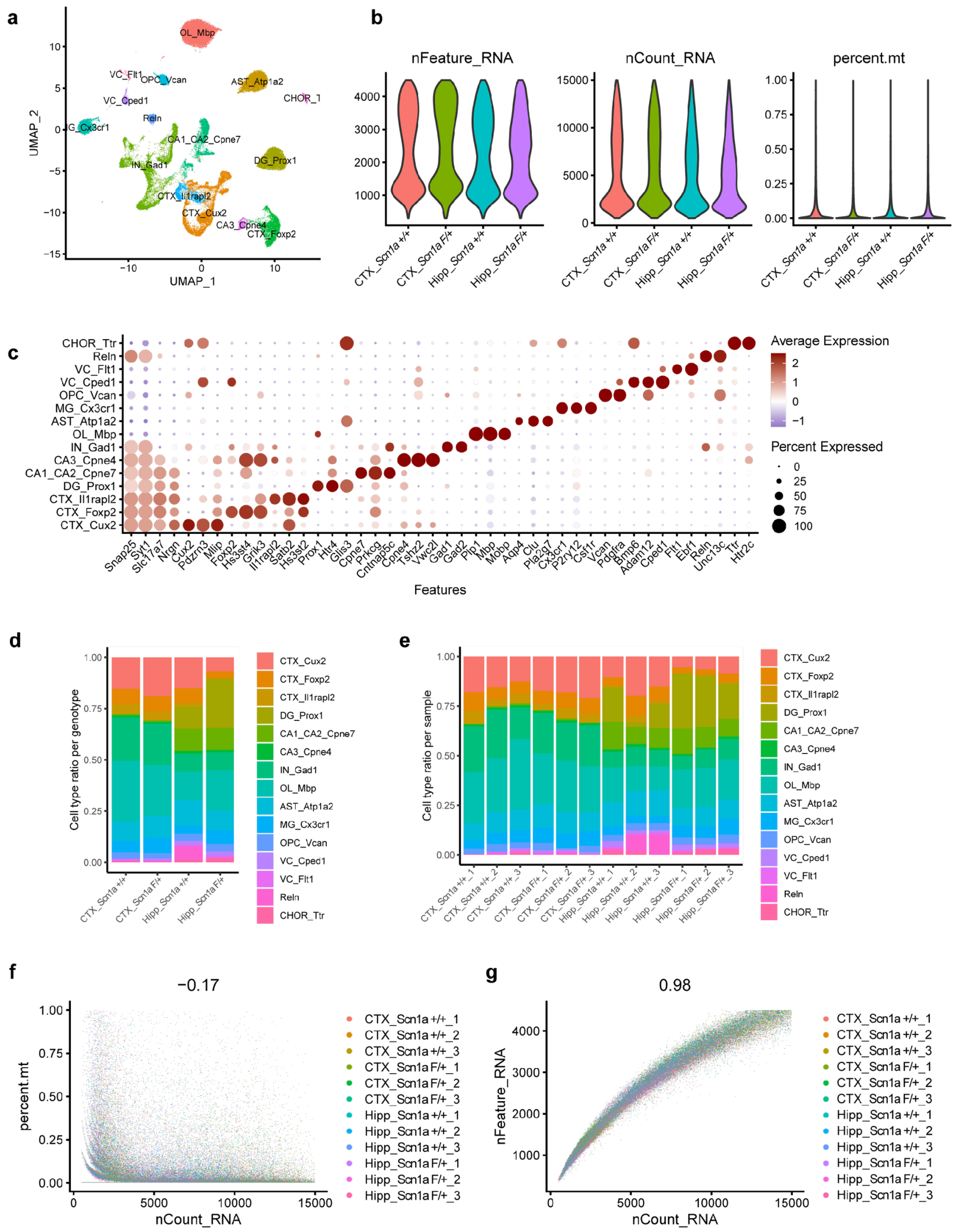


**Extended Data Figure 1. Quality control of the *PV-cre/+* conditional DS snRNAseq dataset (associated with Figure 2)**

(a) UMAP dimensional plot showing transcriptionally distinct cell type clusters identified using Seurat. (b) Violin plots showing equivalent amounts of total number and counts of genes and percent mitochondrial RNA in nuclei used for downstream analyses. (c) Dotplot showing the annotation of cell types based on the average expression and percentage of cells expressing different cell-type specific marker genes in each cluster identified in a). (d) Stacked bar chart showing the ratio of each cell type across brain regions and genotypes of all sequenced nuclei. (e) Stacked bar chart showing the ratio of each cell type in nuclei of each individual sample. (f-g) Correlation between UMI counts and percentage of mitochondrial genes per nuclei (f) and total genes detected (g) for all samples.

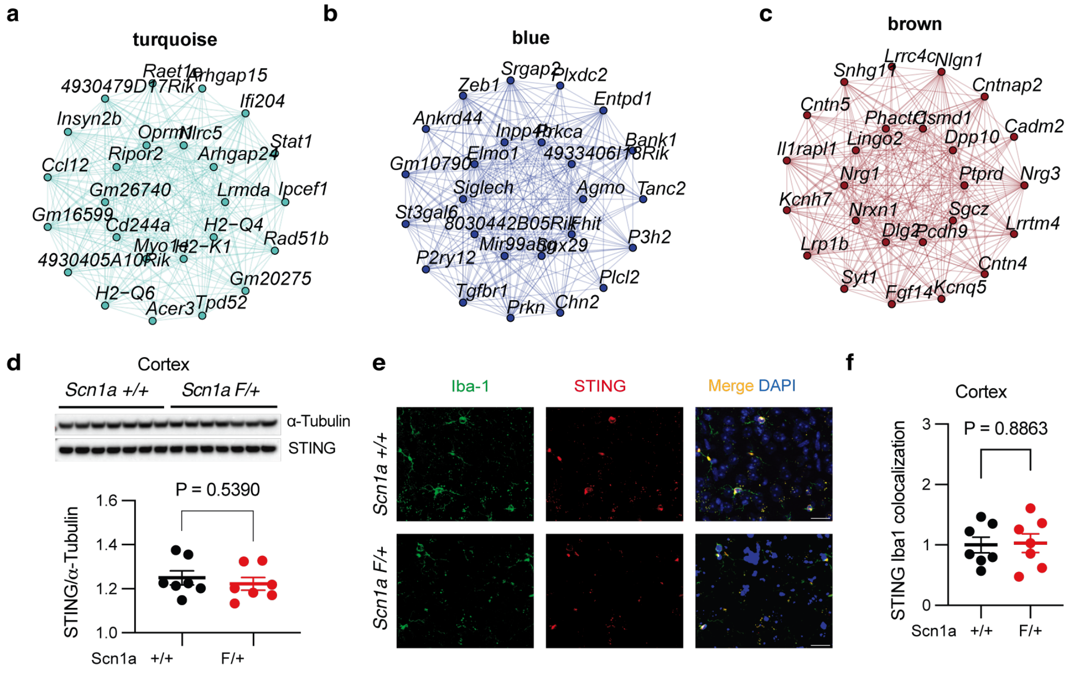


**Extended Data Figure 2. Levels of STING in the cortical tissues.** (a-c) Module network plots visualizing the top 25 hub genes for the turquoise, blue, and brown modules predicted by hdWGCNA in the conditional DS PV-cre/+, Scn1a F/+ microglia. (d) Western blot image and quantification of STING normalized to α-Tubulin in PV-cre/+, Scn1a +/+ mice (n=7) and PV-cre/+, Scn1a F/+ mice (n=7) cortex lysate. Welch's t-test. (e) Representative immunofluorescence images of Iba1 and STING in the cortex of PV-cre/+, Scn1a F/+ mice and control mice. Scale bar, 20µm. (f) Quantification of STING IBA1 colocalization within the cortex. Each circle represents the mean quantification of four brain sections per animal. PV-cre/+, Scn1a +/+ , n=7; PV-cre/+, Scn1a F/+ mice, n=7. Welch's t-test.

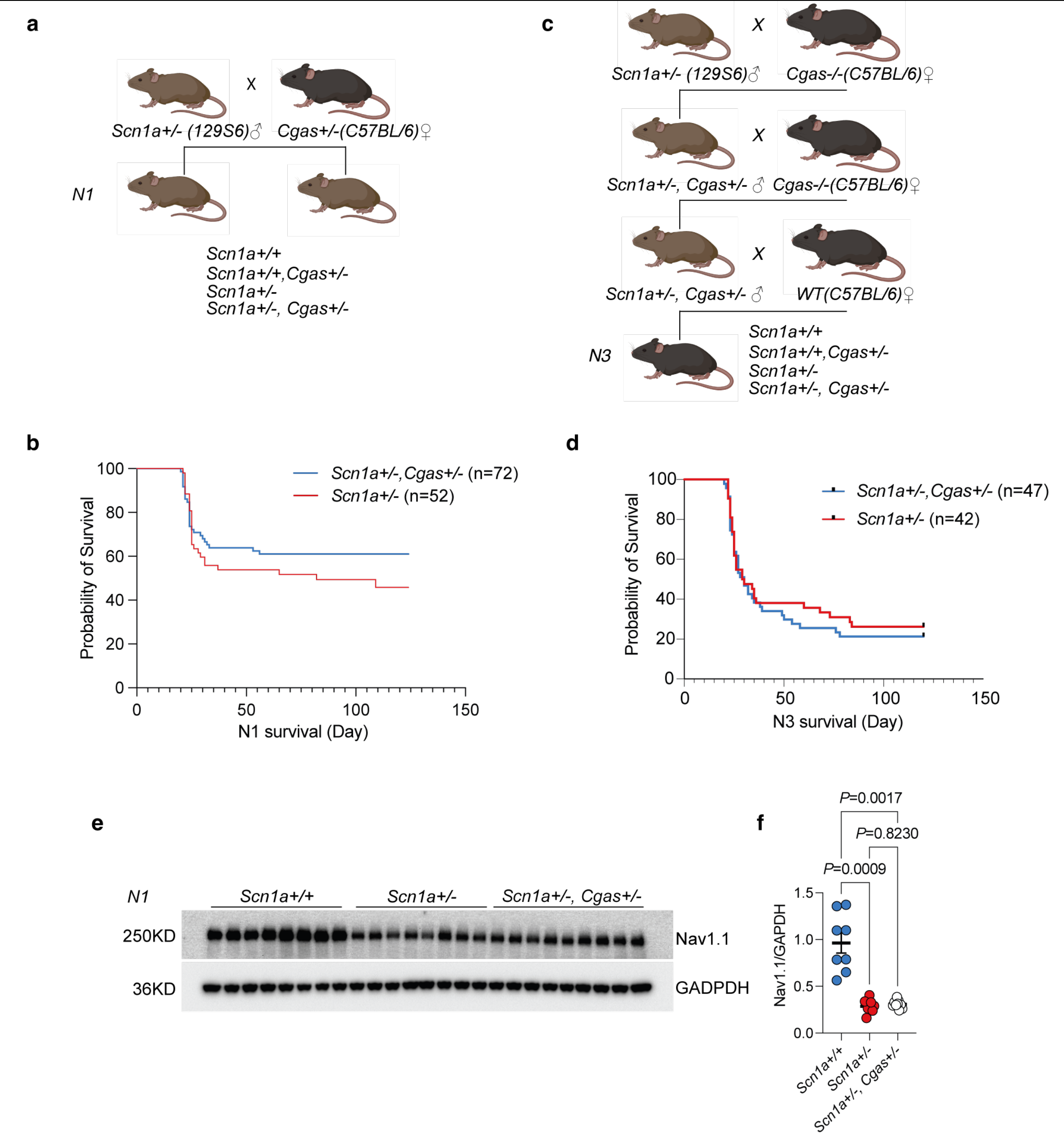


**Extended Data Figure 3. Generation and characterization of the Dravet mouse model (associated with Figure 4).**

(a) Schematic showing the breeding scheme of N1 DS mice. (b) Survival of N1 *Scn1a+/-* (n=52) and *Scn1a+/-, Cgas+/-* (n=72) mice monitored till P120. Log-rank Mantel-Cox test, $P = 0.24$. (c) Schematic showing the breeding scheme of N3 DS mice. (d) Survival of N3 *Scn1a+/-* (n=42) and *Scn1a+/-, Cgas+/-* (n=47) mice monitored till P120. Log-rank Mantel-Cox test, $P = 0.61$. (e) Western blot image and (f) quantification of Nav1.1 protein level normalized to GAPDH in the hippocampal lysate of N1 *Scn1a+/+* (n=8), *Scn1a+/-* (n=8), *Scn1a+/-, Cgas+/-* (n=9) mice. Welch's ANOVA with Dunnett's T3 multiple comparisons test.

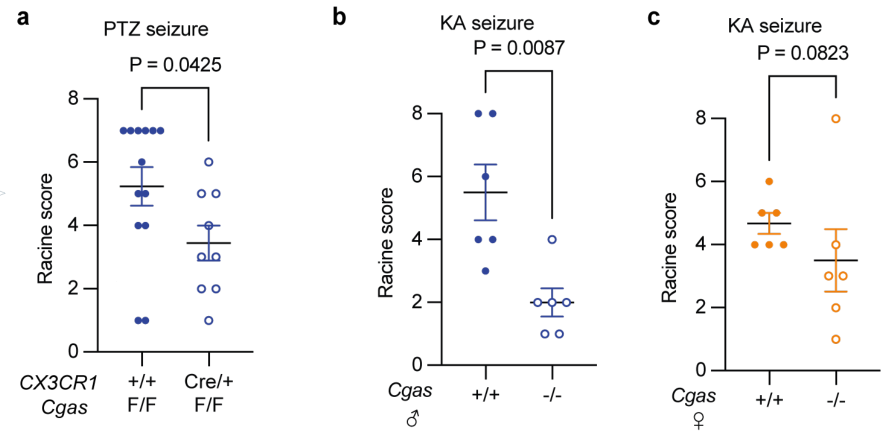


**Extended Data Figure 4. cGAS reduction reduces epileptic events in pharmacological associated seizures (associated with Figure 4).**
(a) Maximum Racine score of seizures induced by PTZ in *cGas-tm1c: F/F; CX3CR1-Cre-ERT2: +/+* mice (n=13) and *cGas-tm1c: F/F; CX3CR1-Cre-ERT2: Cre/+* (n=9). Mann-Whitney test. (b) Maximum Racine score of seizures induced by kainic acid (KA) in male *Cgas+/+* mice (n=6) and *Cgas-/-* mice (n=6). Mann-Whitney test. (c) Maximum Racine score of seizures induced by PTZ in female *Cgas+/+* mice (n=6) and *Cgas-/-* mice (n=6). Mann-Whitney test.

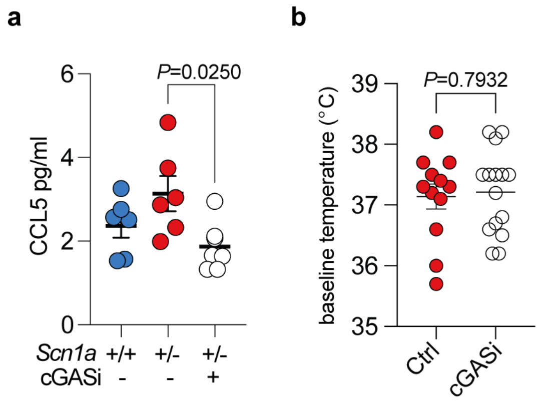


**Extended Data Figure 5. Data related to cGASi treatment (associated with Figure 4)**
(a) Quantification of CCL5 chemokine concentration in the hippocampi lysate from *Scn1a+/+* Ctrl, *Scn1a+/-*, Ctrl, and *Scn1a+/-*, cGASi mice. One-way ANOVA with Tukey's multiple comparisons test. n=6 per genotype. (b) Quantification of the baseline body temperature before hyperthermia-induced seizures in *Scn1a+/-*, Ctrl (n=12), and *Scn1a+/-*, cGASi (n=15) mice. unpaired t test.

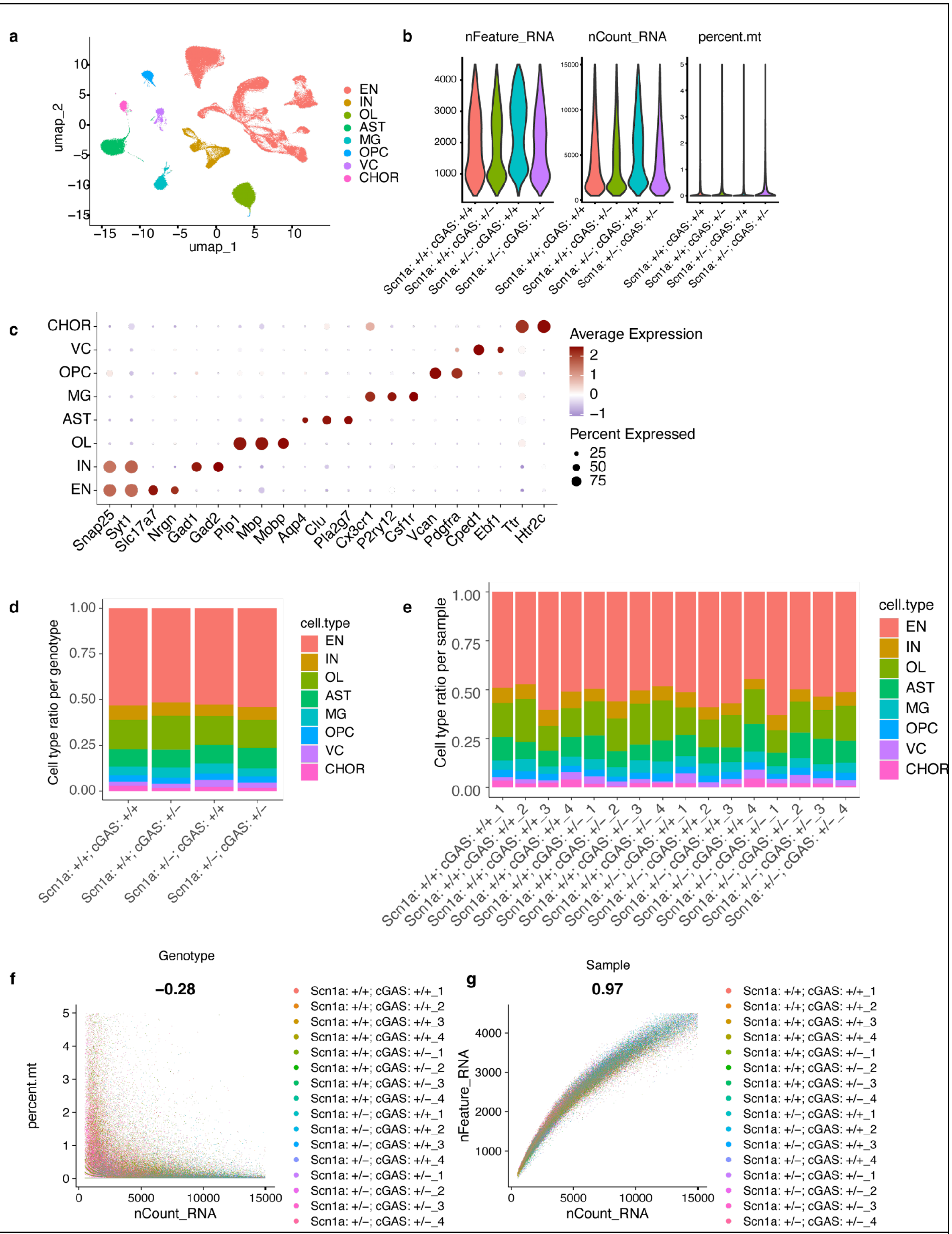


**Extended Data Figure 6. Quality control of the N3 DS snRNAseq dataset (associated with Figure 5, 6)**

(a) UMAP dimensional plot showing transcriptionally distinct cell type clusters identified using Seurat. (b) Violin plots showing equivalent amounts of total number and counts of genes and percent mitochondrial RNA in nuclei used for downstream analyses. (c) Dotplot showing the annotation of cell types based on the average expression and percentage of cells expressing different cell-type specific marker genes in each cluster identified in (a). (d) Stacked bar chart showing the ratio of each cell type across brain regions and genotypes of all sequenced nuclei. (e) Stacked bar chart showing the ratio of each cell type in nuclei of each individual sample. (f,g) Correlation between UMI counts and percentage of mitochondrial genes per nuclei (f) and total genes detected (g) for all samples.

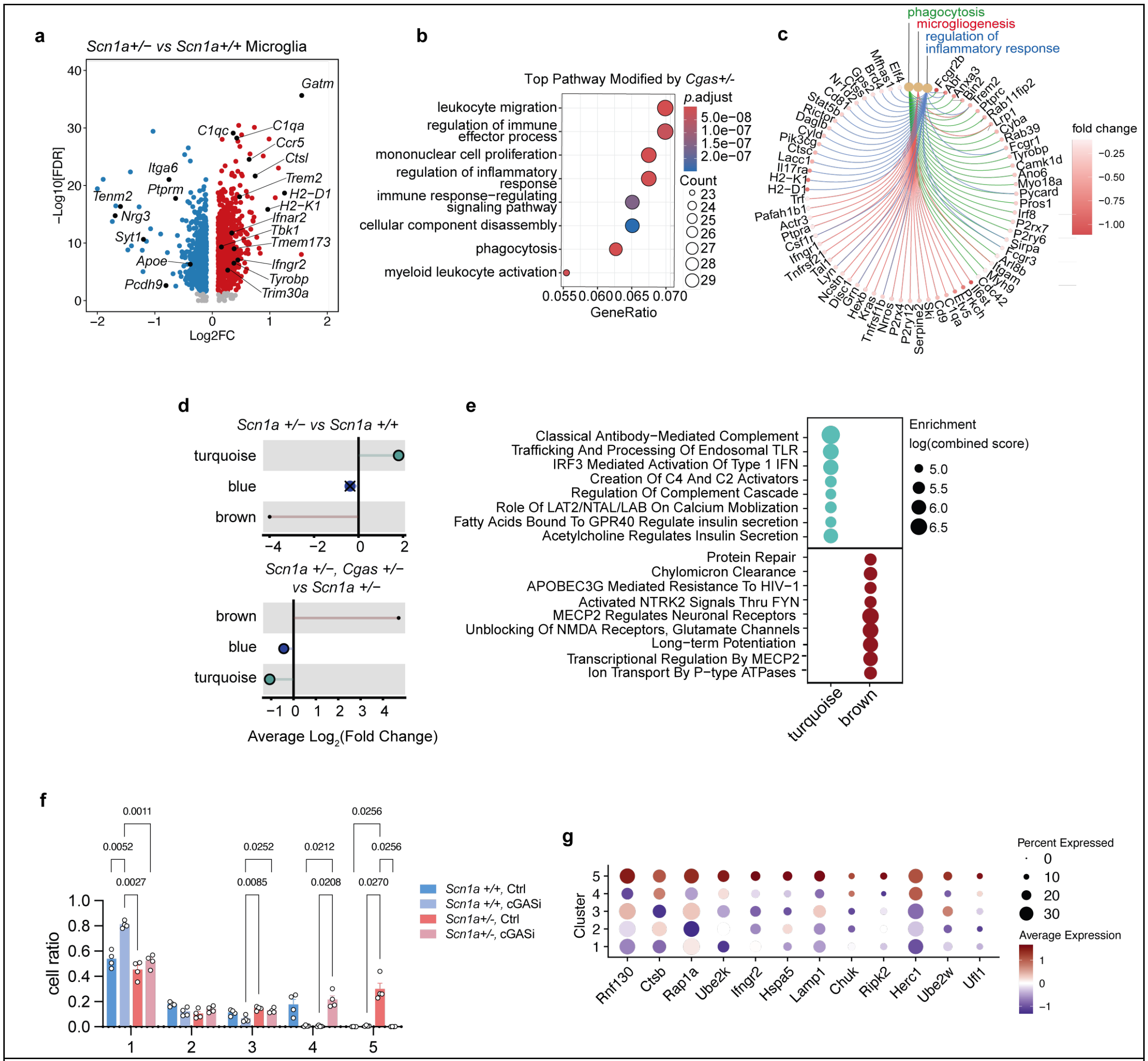


**Extended Data Figure 7. cGAS reduction and inhibition alleviates the inflammatory signature in Dravet microglia (associated with Figure 5).**

(a) Volcano plot of the DEGs between *Scn1a* +/- versus *Scn1a* +/+ microglia. Red and blue dots represent significant DEGs ((logFC >=0.1 or <= -0.1, *p*.adj <0.05). (b) Dotplot showing the top GO pathways downregulated in *Scn1a*+/-, *Cgas*+/- versus *Scn1a*+/- microglia. (c) Cnetplot showing the DEGs associated with selected GO pathways identified in b). (d) Lollipop plot visualizing the hdWGCNA differential module eigengene (DME) analysis results comparing *Scn1a*+/- versus *Scn1a*+/+ (top) and *Scn1a*+/-, *Cgas*+/- versus *Scn1a*+/- microglia (bottom). The size of each dot corresponds to the number of genes in that module. "X" is placed over each point that does not reach statistical significance. (e) Dotplot of the enrichment analysis showing top reactome pathways associated with the marker genes of the turquoise and brown modules. (f) Bar plot of the ratio of microglia that belong to each cluster in each individual mouse by genotype and treatment. Each circle represents an individual animal. Two-way ANOVA analysis with Tukey's multiple comparisons. (g) Dotplot showing the scaled expression level and percentage expression of selected inflammation-related cluster 5 marker genes across microglia clusters.

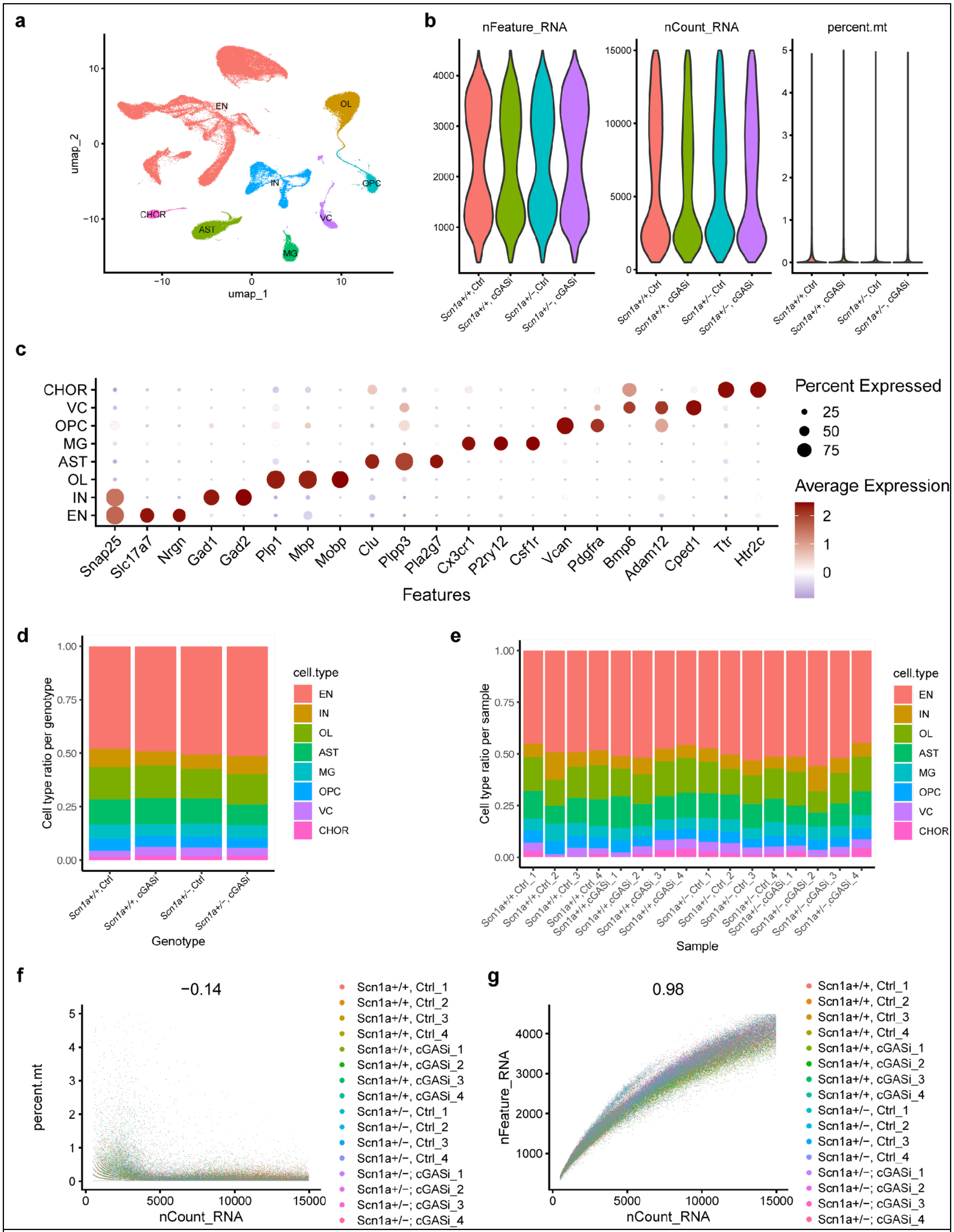

**Extended Data Figure 8. Quality control of the N1 Dravet-cGASi snRNAseq dataset (associated with Figure 5,7)**

(a) UMAP dimensional plot showing transcriptionally distinct cell type clusters identified using Seurat. (b) Violin plots showing equivalent amounts of total number and counts of genes and percent mitochondrial RNA in nuclei used for downstream analyses. (c) Dotplot showing the annotation of cell types based on the average expression and percentage of cells expressing different cell-type specific marker genes in each cluster identified in a). (d) Stacked bar chart showing the ratio of each cell type across brain regions and genotypes in all sequenced nuclei. (e) Stacked bar chart showing the ratio of each cell type in nuclei of each individual sample. (f,g) Correlation between UMI counts and percentage of mitochondrial genes per nuclei (f) and total genes detected (g) for all samples.

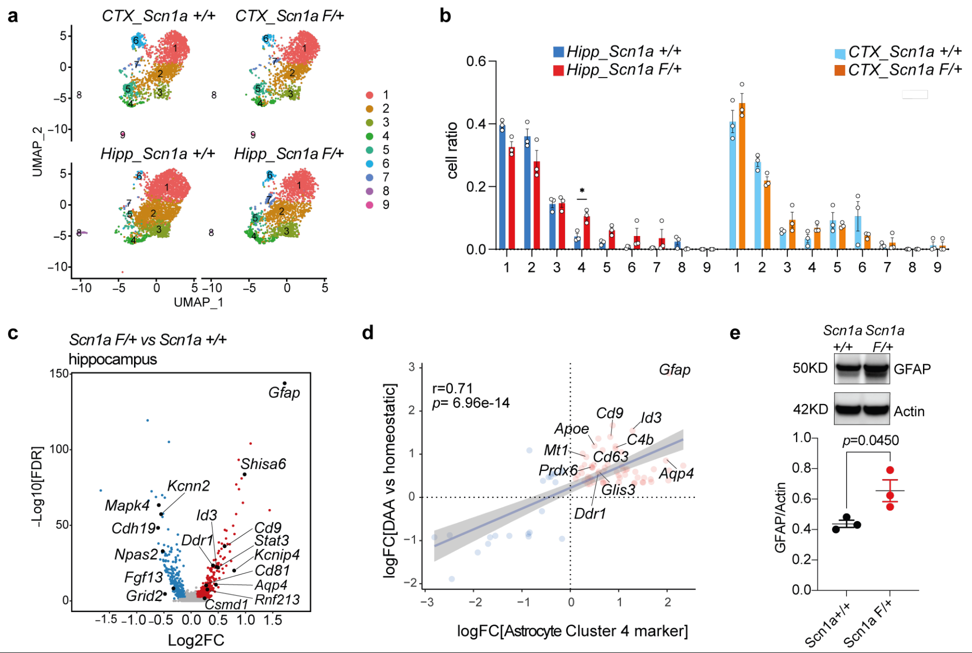


**Extended Data Figure 9. Disease-associated astrocyte signature is observed in the *PV Cre/+* conditional Dravet model.**

(a) UMAP of astrocytes colored by clusters and split by genotype sequenced from the conditional DS mice. (b) Bar plot of the ratio of astrocytes that belong to each cluster in the hippocampus and cortex region of individual *PV-cre/+, Scn1a +/+* and *PV-cre/+, Scn1a F/+* mouse. Each circle represents an individual animal. **P* = 0.0476. Two-way ANOVA analysis with Tukey's multiple comparisons. (c) Volcano plot of the DEGs between *PV-cre/+, Scn1a F/+* versus *PV-cre/+, Scn1a +/+* astrocytes in the hippocampus. Red and blue dots represent significant DEGs ((logFC >=0.1 or <= -0.1, *p*.adj <0.05). (d) Scatter plot and simple linear regression analysis with standard error showing a positive correlation between Cluster 4 astrocyte markers and DAA markers. Pearson's r = 0.71, *p*<0.0001. (e) Representative western blot image and quantification of GFAP protein level normalized to β-Actin in the hippocampal lysate of *PV-cre/+, Scn1a F/+* (n=3) and *PV-cre/+, Scn1a +/+* (n=3) mice. Unpaired t-test.

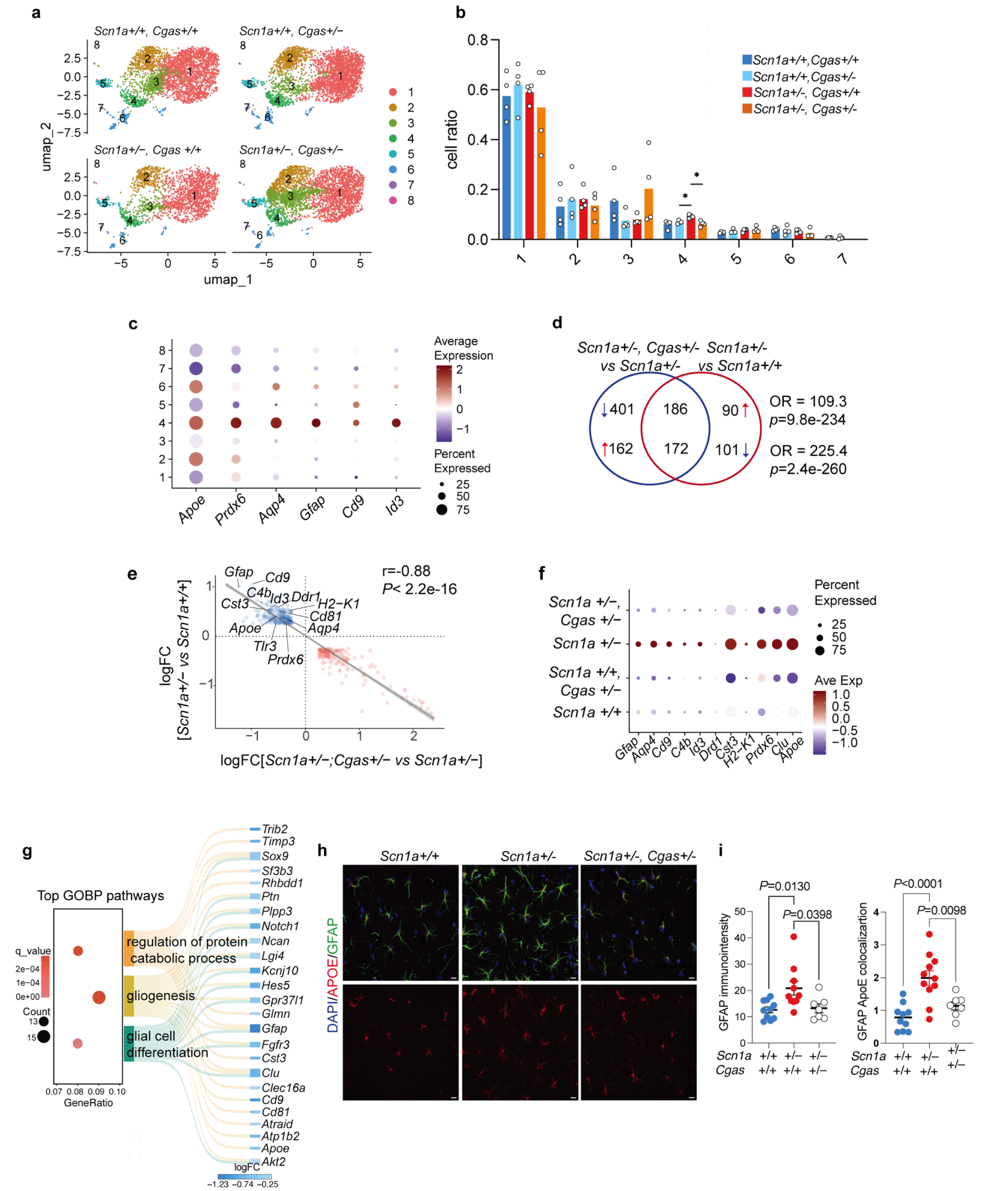


**Extended Data Figure 10. cGAS reduction ameliorates Dravet-induced Disease-associated astrocyte signature.**

(a) UMAP of astrocytes colored by clusters and split by genotype sequenced from the hippocampi of N3 mice. (**b**) Bar plot of the ratio of astrocytes that belong to each cluster in each individual mouse by genotype. Each circle represents an individual animal. *Scn1a+/+, Cgas+/-* vs. *Scn1a+/-, Cgas+/+*, *$P$ = 0.011. *Scn1a+/-, Cgas+/+ vs. Scn1a+/-, Cgas+/-*, *$P$ = 0.024. Two-way ANOVA analysis with Tukey's multiple comparisons. (c) Dotplot showing the expression level and percentage expression of DAA marker genes across astrocyte clusters, suggesting DAA signature is highest in cluster 4. (d) Venn diagram showing the overlap between the DEGs (logFC >=0.25 or <= -0.25, *p*.adj <0.05) of *Scn1a+/-, Cgas+/-* versus *Scn1a+/-*, and *Scn1a+/-* versus *Scn1a +/+* astrocytes. Overlapping odds ratio and *P* value calculated with Fisher's exact test. (e) Scatter plot and simple linear regression analysis with standard error showing a negative correlation between logFC values of overlapping DEGs in D). Pearson's r = -0.88, *p*<2.23e-16. (f) Dotplot showing the scaled expression level and percentage expression of disease-associated astrocyte markers across genotypes with upregulated expression in *Scn1a+/-* astrocytes. (g) Dotplot showing the top pathways upregulated in *Scn1a+/-* and downregulated in *Scn1a+/-, Cgas+/-* astrocytes and Sankey plot showing the DEGs associated with each pathway. (H) Representative 40x immunofluorescence images of GFAP and APOE in the CA1 region of hippocampi of 4–5-month-old N3 mice. Scale bar, 10um. (i) Left: quantification of GFAP immunointensity in the CA1 region of hippocampi of 4–5-month-old N3 mice. Each dot represents the mean intensity of three to four brain sections per animal. One-way ANOVA with Tukey's multiple comparisons test. *Scn1a+/+, Cgas+/+*, n=10; *Scn1a+/-, Cgas+/+*, n=10 (after one significant outlier removed with ROUT test, Q= 1%); *Scn1a+/-, Cgas+/-*, n=7. Right: quantification of APOE immunointensity within GFAP-positive areas. Each dot represents the mean quantification of three to four brain sections per animal. One-way ANOVA with Tukey's multiple comparisons test. *Scn1a+/+, Cgas+/+*, n=10; *Scn1a+/-, Cgas+/+*, n=11; *Scn1a+/-, Cgas+/-*, n=7.

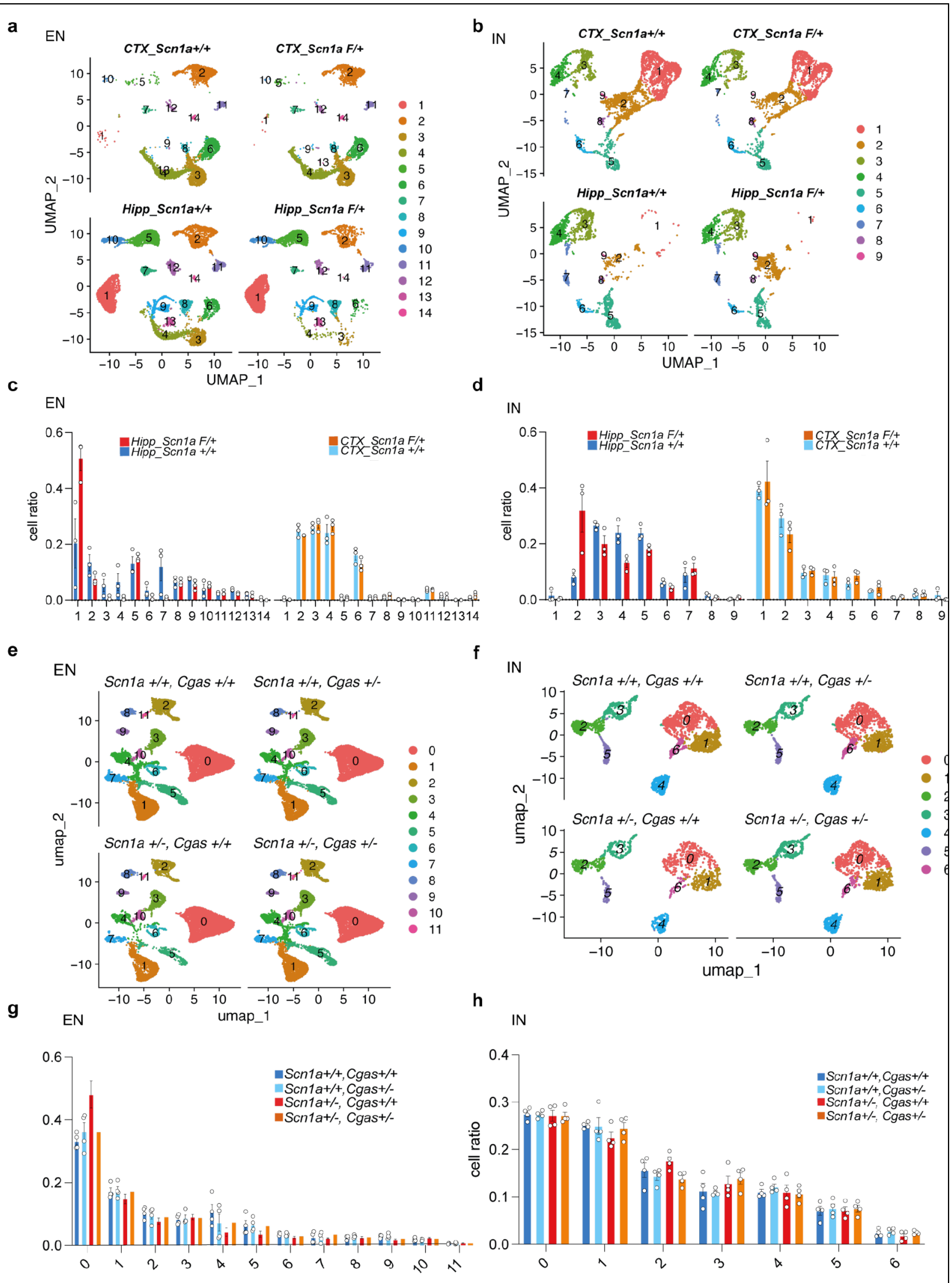


**Extended Data Figure 11. snRNAseq cluster profile for neurons (associated with Figure 6).**

(a, b) UMAP of excitatory neurons (a) and inhibitory neurons (b) colored by clusters and split by genotype and brain regions sequenced from the conditional DS mice.
(c, d) Bar plot of the ratio of excitatory neurons (c) and inhibitory neurons (d) that belong to each cluster in each individual mouse by genotype and brain region. Each circle represents an individual animal. Two-way ANOVA analysis with Sidak's multiple comparisons.
(e, f) UMAP of excitatory neurons (e) and inhibitory neurons (f) colored by clusters and split by genotype sequenced from the hippocampi of N3 DS mice.
(g, h) Bar plot of the ratio of excitatory neurons (g) and inhibitory neurons (h) that belong to each cluster in each individual mouse by genotype. Each circle represents an individual animal. Two-way ANOVA analysis with Tukey's multiple comparisons.

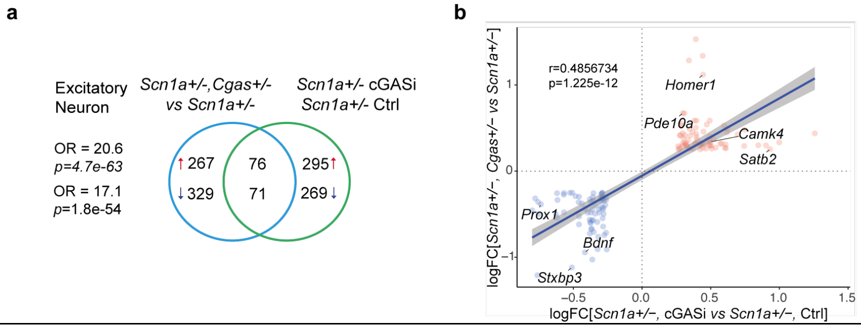


**Extended Data Figure 12. Comparison between neuronal transcriptomic changes associated with cGAS reduction and inhibition (associated with figure 6,7)**
(a) Venn diagram showing the overlap between the top DEGs (logFC >=0.25 or <= -0.25, *p*.adj <0.05) of *Scn1a+/-, Cgas+/-* versus *Scn1a+/-*; and *Scn1a+/-*, cGASi versus *Scn1a +/-*, Ctrl excitatory neurons. Overlapping odds ratio and *P* value calculated with Fisher's exact test. (b) Scatter plot and simple linear regression analysis with standard error showing a positive correlation between logFC values of overlapping DEGs in A). Pearson's r = 0.49, *P*<0.0001.

**Supplemental tables**

**Table S1. scRNAseq data of the DRE microglia reported by Kumar et al, 2022.** Marker genes for DRE microglia clusters. Selected top transcription factors controlling the expression of MG4 marker genes. Selected GSEA hallmark pathways associated with microglia cluster 4 markers. Significant IPA upstream regulators predicted for microglia cluster 4 (Activation Z score >=2 or <=-2, *p*-value of overlap <0.05)

**Table S2. snRNAseq data of the TLE microglia reported by Chen et al, 2023.** DEGs between microglia of TLE versus Control participants. Selected activated IPA upstream regulators predicted based on microglia DEGs. Selected top GSEA hallmark pathways associated with microglia DEGs.

**Table S3 Human epilepsy patient data associated with IHC stainings in Fig 1.**

**Table S4 snRNAseq data of the conditional DS model.** DEG in hippocampal microglia of *PV-Cre/+, Scn1a F/+* vs *PV-Cre/+, Scn1a +/+* mice. Marker genes of each hdWGCNA module in microglia. Marker genes of each astrocyte cluster. DEG in hippocampal astrocyte of *PV-Cre/+, Scn1a F/+* vs *PV-Cre/+, Scn1a* +/+ mice.

**Table S5 snRNAseq data of the N3 cGAS-DS model.** DEGs between *Scn1a+/- vs Scn1a* +/+ hippocampal microglia. DEGs between *Scn1a+/-, Cgas+/- vs Scn1a* +/- hippocampal microglia. Enrichment analysis of the Reactome pathways associated with microglia hdWGCNA modules. DEGs between *Scn1a+/- vs Scn1a* +/+ hippocampal astrocytes. DEGs between *Scn1a+/-, Cgas+/- vs Scn1a* +/- hippocampal astrocytes. DEGs between *Scn1a+/- vs Scn1a* +/+ hippocampal excitatory neurons. DEGs between *Scn1a+/-, Cgas+/- vs Scn1a* +/- hippocampal excitatory neurons. DEGs between *Scn1a+/- vs Scn1a* +/+ hippocampal inhibitory neurons. DEGs between *Scn1a+/-, Cgas+/- vs Scn1a* +/- hippocampal inhibitory neurons.

**Table S6 snRNAseq data of the N1 cGASi-DS model.** Marker genes for microglia clusters. DEGs between *Scn1a+/-* Ctrl *vs Scn1a* +/+ Ctrl hippocampal microglia. DEGs between *Scn1a+/-,* cGASi *vs Scn1a* +/- Ctrl hippocampal microglia. DEGs between *Scn1a+/-* Ctrl *vs Scn1a* +/+ Ctrl hippocampal excitatory neurons. DEGs between *Scn1a+/-,* cGASi *vs Scn1a* +/- Ctrl hippocampal excitatory neurons. DEGs between *Scn1a+/-* Ctrl *vs Scn1a* +/+ Ctrl hippocampal inhibitory neurons. DEGs between *Scn1a+/-,* cGASi *vs Scn1a* +/- Ctrl hippocampal inhibitory neurons.
